# The DREAM complex links somatic mutation, lifespan, and disease

**DOI:** 10.1101/2025.09.15.676396

**Authors:** Zane Koch, Shuvro P. Nandi, Kate Licon, Arturo Bujarrabal-Dueso, David H. Meyer, Safa Saeed, Pirunthan Perampalam, Frederick A. Dick, Björn Schumacher, Ludmil B. Alexandrov, Trey Ideker

## Abstract

The DREAM complex has emerged as a central repressor of DNA repair, raising questions as to whether such repression exerts long-term effects on human health. Here we establish that DREAM activity significantly impacts lifetime somatic mutation burden, and that such effects are linked to altered lifespan and age-related disease pathology. First, joint profiling of DREAM activity and somatic mutations across a single-cell atlas of 21 mouse tissues shows that cellular niches with lower DREAM activity have decreased mutation rates. Second, DREAM activity predicts the varied lifespans observed across 92 mammals, with low activity marking longer-lived species. Third, reduced DREAM activity in Alzheimer’s patients predicts late disease onset and decreased risk for severe neuropathology. Finally, we show DREAM knockout protects against mutation accumulation *in vivo*, reducing single-base substitutions by 4.2% and insertion/deletions by 19.6% in brains of mice. These findings position DREAM as a key regulator of aging.

## Introduction

To protect against the constant barrage of DNA lesions arising from endogenous cellular processes and exogenous damaging agents, organisms have evolved an extensive network of DNA repair mechanisms^1^. These mechanisms are especially effective in long-lived species, which exhibit increased resistance to cellular stresses^2,3^, decreased somatic mutation rates^4,5^, and more efficient DNA repair^6^. Elevated DNA repair activity is also present in reproductive (germline) tissues, yielding mutation rates that are up to 100-fold lower than in the soma^7–9^. Genomic studies in centenarians have found strong enrichments for specific genetic variants that activate DNA repair genes^10–12^; conversely, genomic instability is a common feature of nearly all progeroid diseases^13,14^ and has been reported to engender numerous phenotypes of old age including neuroinflammation^15^, type 2 diabetes^16^, kidney dysfunction^17,18^, and Alzheimer’s disease (AD)^19–22^. This prioritization of DNA repair in long-lived species, reproductive tissues, and centenarians underscores the importance of genomic integrity to longevity and raises questions about how such differences in DNA repair arise.

Recently, a protein complex known as DREAM has been shown to transcriptionally repress DNA repair pathways in somatic cells, where these pathways are not necessary for reproductive success^23^. The DREAM complex (named for its subcomponents, <u>Dp</u>, <u>R</u>b-like-1, <u>E</u>2f, <u>A</u>nd <u>M</u>uvB) was originally characterized as a repressor of the cell cycle^24^. It forms as cells enter quiescence (G0 phase), when MuvB is phosphorylated by the dual-specificity tyrosine-regulated kinase 1A (DYRK1A)^25^, leading to DREAM formation and the repression of numerous target genes^26,27^. To later exit quiescence, the components of the DREAM complex are prompted to dissociate by the phosphorylation of p130 (*RBL2*) by cyclin-dependent kinases^24,28^, lifting repression of DREAM targets^26^. Following these cell-cycle studies, the direct targets of the DREAM complex have been shown to include many genes functioning in DNA damage response (DDR)^23,29^. Disruption of DREAM in nematode worms, progeroid mice, or human cells produces resistance to DNA damaging agents^23^, whereas cells with an extra copy of DYRK1A – and thus greater DREAM activity^30^ – show increased levels of γH2AX, a marker of DNA damage^31^.

Collectively, these recent results have extended the role of DREAM from repression of cell cycle to repression of DDR and its associated functions. Here we seek to determine if the DREAM complex not only governs the cell cycle and DNA repair but also hallmarks of aging and longevity (**Fig. 1a**). We find that, at the cellular level, DREAM activity associates with somatic mutation rate; across species, DREAM activity inversely correlates with maximum lifespan; in humans, diminished DREAM activity predicts later and less severe Alzheimer’s disease pathology; and in mice, induced DREAM deficiency slows the rate of somatic mutation.

**Figure 1:**
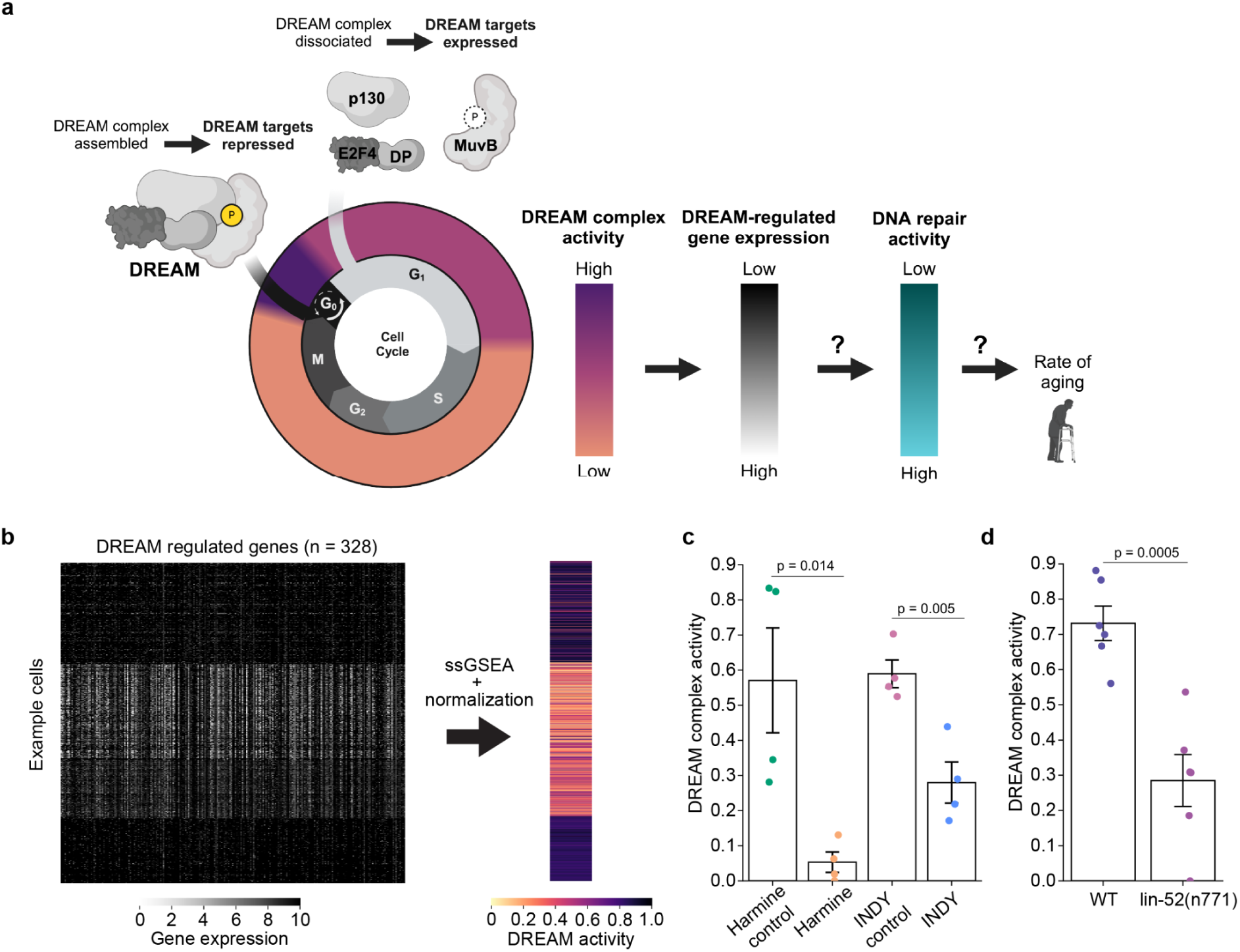
Transcriptional measurement of DREAM complex activity. **a)** The DREAM transcriptional repressor complex is formed in G0 following the phosphorylation of MuvB by DYRK1A. Upon reentry to the cell cycle, the DREAM complex dissociates and DREAM-target genes can be expressed. Here we explore the relationship between DREAM activity and somatic mutation, as well as between DREAM activity and rate of aging. **b)** Calculation of DREAM activity from target gene expression. Single sample gene set enrichment analysis (ssGSEA) is used to summarize the mRNA expression values of all detected DREAM complex target genes (columns, 328 DREAM-regulated genes) in each cell (rows). This ssGSEA score is then normalized and corrected for sequencing depth to produce a per cell DREAM activity score (**Methods**). **c)** U2OS cells are exposed to a DREAM complex inhibitor (harmine, INDY) versus control treatment (harmine control, INDY control). mRNA expression profiles are measured, from which we compute DREAM activity. Bar heights indicate mean DREAM activity across samples (n = 4 biological replicates), error bars denote 95% confidence intervals, and p-values indicate the result of two-sided t-tests. **d)** Similar to (c), but showing DREAM complex activity in wild type (WT) vs. *lin-52*(*n771*) mutant *C. elegans*.

## Results

### Transcriptional readout of DREAM repressor activity

We selected genes with regulatory regions previously shown to be bound by multiple components of the DREAM complex through protein-DNA interactions^23,24^ (n = 328 genes). The expression of these DREAM-target genes was used to score DREAM complex activity (**Methods**), signed such that low expression corresponds to high DREAM activity (**Fig. 1a-b**).

The utility of this DREAM activity score was underscored by several observations. First, in human tissues this score was tightly associated with the protein abundance and modification status of the DREAM-subunits MuvB and p107/p130, whose phosphorylation status controls DREAM assembly^28^ (Pearson r = 0.70, p = 6.8 × 10^−15^, n = 160 human individuals, **Supplementary Fig. 1a-b, Methods**). Second, the score successfully distinguished cells treated with DREAM complex inhibitors^23^, harmine or INDY, from those of control cells (**Fig. 1c**). Third, gene expression analysis of nematode worms with a mutation in *lin-52*, which causes defects in DREAM complex assembly, yielded an activity score that was markedly lower than that of wild-type worms^23^ (**Fig. 1d**). Fourth, increased DREAM activity was associated with lower expression of the proliferative marker MKI67^32^ across cell types (Spearman ρ = –0.56, p = 4.37 × 10^−11^, n = 120 cell types, **Supplementary Fig. 1c**) and tissues (Spearman ρ = –0.45, p = 0.03, n = 23 tissues, **Supplementary Fig. 1d**) – as would be expected given the DREAM complex’s role as a negative regulator of the cell cycle^24,26^. Collectively, these results suggested that the DREAM activity score is an informative and specific metric for quantifying the repressive activity of the DREAM complex.

### DREAM activity associates with somatic mutation rate in single cells

We used this DREAM activity measure to study the connection between DREAM and somatic mutation. For this purpose we analyzed the *Tabula Muris Senis* dataset (TMS, **Supplementary Table 1**)^33^, which includes full-length RNA sequencing^34^ of single cells collected from 21 organs in 18 C57BL/6JN mice at young (3 months), middle (18 months), and old (24 months) age. DREAM activity was scored in the single-cell RNA-seq data of each of 110,824 cells, revealing a broad range of values that varied significantly across organs and individual mice, and decreased with age (**Supplementary Fig. 2a-c**).

Somatic mutations were called from the same scRNA-seq data, leveraging a previous analysis^33^ (**Methods**). Here we observed that the overall burden of somatic mutations increased with age in nearly every tissue (**Fig. 2a**), corresponding to rates of mutation ranging from a low of 0.0025 mutations/kb/month (tongue) to a high of 0.0045 (kidney).

**Figure 2:**
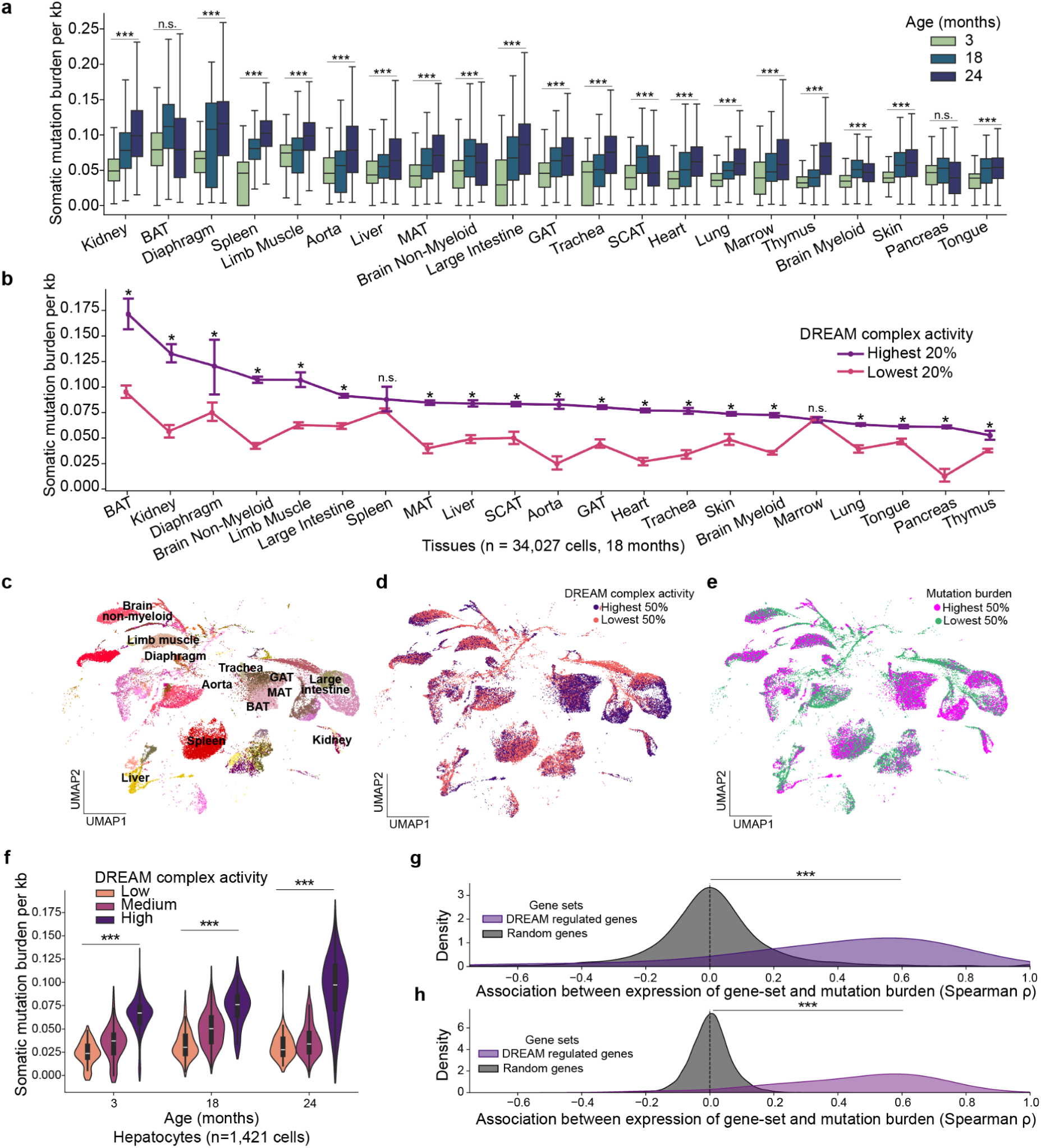
Association between DREAM activity and single-cell somatic mutation rate. **a)** Box plots of the distribution of somatic mutation burden across the cells of each tissue at each age (n = 110,824 cells, n = 18 mice). Mutation burden is expressed as the mutation count per callable kilobase (**Methods**). (***) and (n.s.) indicate p values calculated by modeling Spearman ρ’s as a Student’s t distribution, yielding p < 5.0 × 10^-19^ and p ≥ 0.01, respectively. BAT, GAT, MAT, and SCAT abbreviate brown, gonadal, mesenteric, and subcutaneous adipose tissue, respectively. Boxes show inter-quartile range (IQR) with median line; whiskers extend to 1.5 IQR. **b)** The somatic mutation burden per kilobase in cells from 18 month old mice (n = 34,027 cells, n = 4 mice) stratified by DREAM complex activity (highest vs. lowest 20% of DREAM complex activities within each tissue). Error bars denote 95% confidence intervals. (*) indicates p < 0.0023 (Bonferroni corrected p-value) and (n.s.) indicates p ≥ 0.0023, based on a two-sided Mann-Whitney test. **c)** Uniform manifold approximation and projection (UMAP) of all cells of all age mice from the first 12 tissue types in (b) (from the left, n = 42,508 cells, n = 18 mice), colored by cell type and labeled with tissue of origin. **d)** As in (c) but coloring cells by DREAM activity. Cells with the 50% lowest and 50% highest DREAM activities for age and tissue type are colored orange and purple, respectively. **e)** As in (d) but coloring cells by somatic mutation burden per kb. Cells with the 50% lowest and 50% highest somatic mutation burdens for their age and tissue type are colored green and pink, respectively. **f)** Violin plot of the somatic mutation burden in hepatocyte cells (n = 1,162 cells, n = 18 mice) stratified by age and DREAM complex activity. (***) indicates a significant (p < 5.0 × 10^-200^) Spearman correlation between DREAM complex activity and mutation burden within an age group. Spearman ρ: 3 months = 0.16, 18 months = 0.30, and 24 months = 0.21. **g)** Kernel density plot depicting the distribution of Spearman ρ’s computed in each cell type (n = 116 cell types, n = 34,027 cells from 18 month old mice, n = 4 mice) between somatic mutation burden and gene-set activity scores (ssGSEA, **Methods**). Purple: DREAM-regulated gene set (n = 248 genes). Gray: 500 random gene sets (n = 248 random genes per set). (***) indicates a significant (p < 4.86 × 10^-14^) difference in Spearman correlation values between groups based on a two-sided Mann Whitney U test. **h)** Similar to (g), but comparing somatic mutation burden and gene expression within *tissue* types (n = 21 tissue types, n = 34,027 cells from 18 month old mice, n = 4 mice, **Methods**).

Notably, in the vast majority of tissues, cells with high DREAM activity had significantly greater mutation burdens than those exhibiting low activity (**Fig. 2b-e**, **Supplementary Fig. 2d-e**). For example, kidney cells in the top quintile of DREAM activity, showed a 2.4-fold increase in somatic mutations per kb compared to cells in the bottom quintile (p = 3.3 × 10^−49^, n = 180 cells per quintile, two-sided Mann-Whitney U test). This positive association between DREAM activity and mutation burden was observed not only at the tissue level but also within individual cell types within a tissue. For instance, liver hepatocytes with greater DREAM activity had significantly greater mutation burdens at each age (**Fig. 2f**), and this was the case for the majority of cell types (**Fig. 2g**) and tissues (**Fig. 2h**) even when correcting for cell proliferation rate (**Supplementary Fig. 2f-g**). In contrast to DREAM target genes, the expression levels of randomly chosen gene sets were not significantly associated with mutation burden (cell types: p = 2.6 × 10^−32^, tissue types: p = 4.86 × 10^−14^; **Fig. 2g-h**, **Methods**).

### DREAM activity predicts lifespan and mutation rate across 92 species

To relate these results to lifespan, we scored DREAM activity in the transcriptomes of 92 mammalian species, broadly spanning six taxonomic orders^35^ (∼3 individuals ⨉ 3 tissues per species, n = 803 samples, **Methods**, **Fig. 3a**). Comparing DREAM activity scores to the maximum lifespan of each species, we observed a strong inverse relationship (**Fig. 3b**). Negative associations were observed across all tissues profiled (**Fig. 3c**) and became even stronger after controlling for the adult weight (AW) of each species (Spearman ρ: liver = –0.59, p = 6.23 × 10^−24^; brain = –0.46, p = 1.26 × 10^−13^; kidney = –0.30, p = 6.23 × 10^−06^; **Fig. 3d**, **Supplementary Fig. 3a-b**). Furthermore, DREAM activity levels tended to be coordinated across all tissues within a species (**Fig. 3e**).

**Figure 3:**
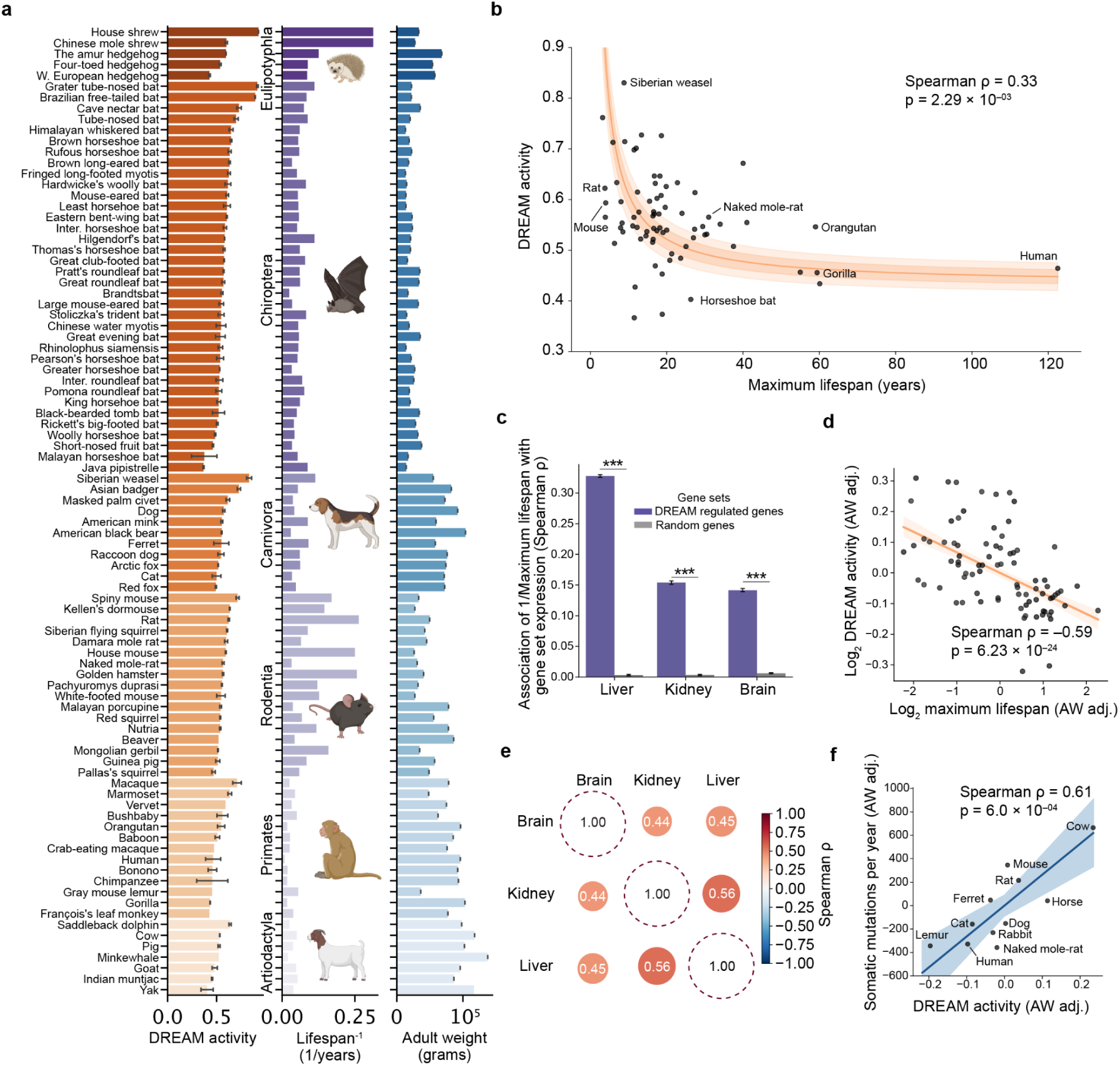
Inverse relation between DREAM activity and lifespan across species. **a)** Barplots indicating the average DREAM activity across tissues (liver, kidney, brain), 1/maximum lifespan, and log_10_ adult weight in grams for each species (n = 92 species and 803 samples). Within each taxonomic order, species are sorted by decreasing average DREAM complex activity. **b)** Scatterplot of maximum lifespan vs. DREAM activity measured in the liver (n = 92 species and 241 samples, **Methods**). Points indicate the mean value of each species, while the orange curve shows reduced major axis regression line fit to the actual values of all samples. The inner shaded area indicates one standard deviation from the regression line and the outer the 95% confidence interval. **c)** Barplots indicating the mean and 99% confidence interval of the Spearman correlation between 1/lifespan and average DREAM activity of each species, in each tissue, iteratively leaving one species out (n = 92 species). (***) indicates p < 8.23 × 10^-24^ based on a two-sided Mann-Whitney test. **d)** Scatterplot and reduced major axis regression depicting the average adult weight (AW) adjusted DREAM activity and lifespan of the species in (a) (n = 92 species and 241 liver samples, **Methods**). Both axes are shown on a logarithmic scale. **e)** Heatmap of the Spearman correlation coefficients comparing DREAM activity scores between each pair of tissues, across the species and samples in (a). All pairwise correlations are significant (p < 0.0005). **f)** Scatterplot and reduced major axis regression depicting the average AW adjusted DREAM activity and somatic mutation rate (in the liver and colon, respectively) across species (n = 11 species, **Methods**).

Comparing the DREAM activity of liver, brain, and kidney tissues to previous measurements of species-specific somatic mutation rates^4^ (n = 11 species), we found that DREAM complex activity in the liver and brain was positively correlated with somatic mutation rate across species (Liver: ρ = 0.61, p = 6.0 × 10^−4^; Brain: ρ = 0.49, p = 0.008; Kidney: ρ = –0.31, p = 0.12; **Fig. 3f, Supplementary Fig. 3c-g, Methods**). Further linking DREAM complex activity to species-specific DNA repair capacity, we observed that fibroblasts from species with lower DREAM complex activity exhibited greater resistance to various stress-inducing agents including the mitochondrial respiration inhibitor rotenone; the oxidative damage inducing agents cadmium and paraquat; the alkylating agent methyl methanesulfonate (MMS); UV radiation; and heat (n = 37 cell lines from 6 species^2^, **Supplementary Fig. 4a-h, Methods**). These results indicated that the amount of DREAM complex activity within a species is associated with the lifespan of that species, as well as its somatic mutation rate and resistance to DNA damage.

### Low DREAM activity protects survival on a high-fat diet

A high-fat diet has been shown to shorten lifespan and induce genomic instability in multiple species^36–38^, associated with an increased burden of oxidative damage and rate of somatic mutation^39^. Using a large cohort of mice from 50 distinct inbred strains^40^ (n = 882 mice, **Supplementary Fig. 5a**), half fed a high-fat diet (HFD; 60% calories from fat) and half fed a regular chow diet (CD; 6% calories from fat), we investigated whether strains with lower DREAM activity were protected from the harms of HFD leading to extended survival.

For each strain and diet, 70% of mice (n = 612 mice) were followed to measure lifespan, while the remaining 30% were sacrificed and transcriptionally profiled to score DREAM activity (n = 270 mouse livers). Lifespan and DREAM activity varied greatly across strains (median lifespan: 704 ± 147 days, mean ± std; **Methods**), with HFD mice having markedly shorter lifespans than CD mice (median survival: HFD = 640 days, CD = 730 days; **Supplementary Fig. 5b-f**). Notably, strains with lower DREAM activity had extended survival on the HFD compared to those with higher DREAM activity (median survival: lowest 20% DREAM activity = 678 days, highest 20% DREAM activity = 578 days; **Fig. 4a-b**). Controlling for sequencing depth and physiological factors – including weight, weight gain on diet, and blood lipid levels (**Supplementary Fig. 5g**, **Methods**) – a one standard deviation increase in DREAM activity was associated with a 36% increase in risk of death for HFD mice (Hazard ratio = 1.36, p = 9.0 x 10^-4^; **Fig. 4b**). When fed the CD, there was no significant difference in survival associated with DREAM activity (Hazard ratio = 0.93, p = 0.27; **Fig. 4b**, **Supplementary Fig. 5h**).

**Figure 4:**
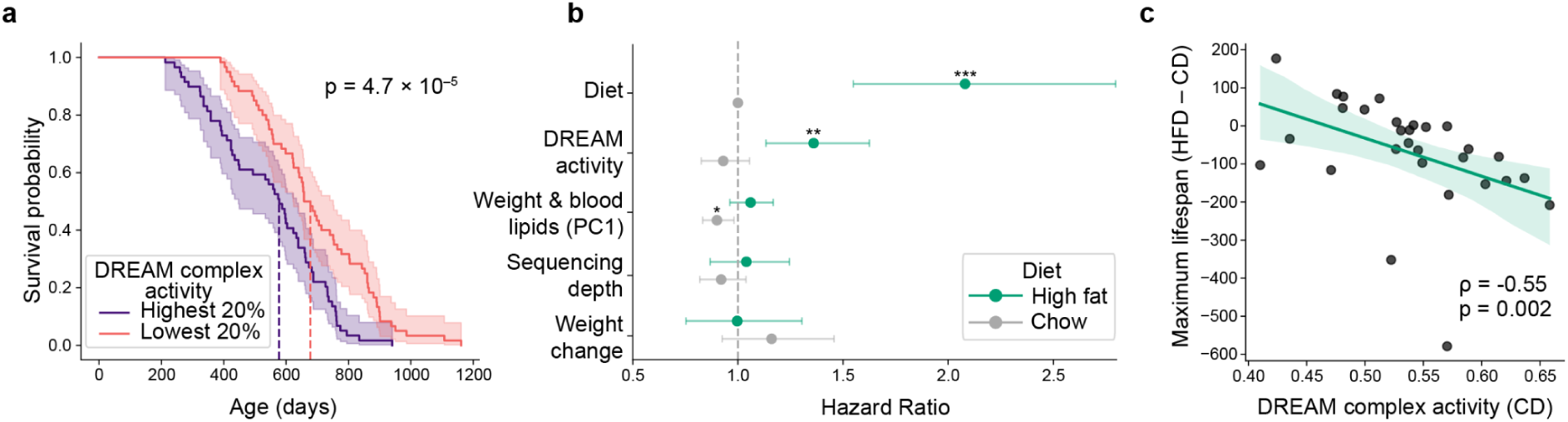
Lifespan of mice with low versus high DREAM activity. **a)** Kaplan-Meier survival curve of high fat diet (HFD)-fed mice belonging to strains with the highest vs. lowest 20% of DREAM complex activities (purple vs. pink, n = 119 mice, n = 14 strains). The dark line and shaded area indicate the proportion of mice surviving to a particular age and the 95% confidence interval of this survival estimate, respectively. Dashed lines indicate the median survival of each group. P value calculated using a two-sided log-rank test. **b)** Dot-and-whisker plot indicating the association of each variable on the y-axis with mouse survival, for mice fed the HFD (green, n = 299 mice) or chow diet (CD) (grey, n = 313 mice). Dots and whiskers indicate the hazard ratio and 95% confidence intervals from a Cox proportional hazard model (**Methods**). (***), (**), and (*) indicate p values, from a two-sided Wald test, of less than 5.0 × 10^-6^, 0.005, and 0.05, respectively. **c)** Scatterplot of mean DREAM activity of each mouse strain on the CD (x axis), compared to the change in survival when members of that strain are placed on the HFD. n = 29 strains profiled on both diets; n = 10 mice per diet per strain on average. P value calculated by modeling Spearman ρ’s as a Student’s t distribution.

We considered whether the association between DREAM activity and lifespan in the HFD mice could be confounded by effects of the high-fat diet itself, whereby the diet increased DREAM activity while decreasing lifespan. However, this interpretation was contradicted by the observation that the DREAM complex activity of a strain when fed the CD was indicative of the survival of mice of the same strain when fed the HFD (Spearman ρ = –0.55, p = 0.002; **Fig. 4c**); that is, higher CD DREAM activity predicted shortened survival when challenged by the HFD. Collectively, these findings suggested that inherent differences in DREAM activity determine resilience to HFD-induced stress, with lower DREAM activity conferring a survival advantage under metabolic challenge.

### DREAM activity is linked to neuropathy and Alzheimer’s disease

We next moved from mice to human samples to test the relevance of our results to diseases associated with human aging. For this purpose, we applied the SComatic^41^ method to identify somatic single nucleotide substitution mutations in 803,122 single cells profiled using single-cell RNA sequencing by the Tabula Sapiens^42^ and Seattle Alzheimer’s Disease Brain Cell Atlas (SEA-AD^43^) consortia (**Fig. 5a**, **Methods**). Cells had been drawn from 11 tissues of 90 human individuals, including 80 with Alzheimer’s Disease and 10 without disease, with donors ranging from 33 to 102 years of age (85.2 ± 13.5 years, mean ± std). Somatic mutations accumulated in tissues at varying rates: from approximately 5 per cell per year in bladder and lung cells, to 10 in brain cells (**Fig. 5b**), and 22 in fat cells (**Fig. 5c, Supplementary Fig. 6a-f**). Notably, we observed that human cells with the highest DREAM activity in each tissue obtained somatic mutations at substantially greater rates than the cells with the lowest DREAM activity (**Fig. 5d**), replicating the results we found in mice (**Fig. 2**).

**Figure 5:**
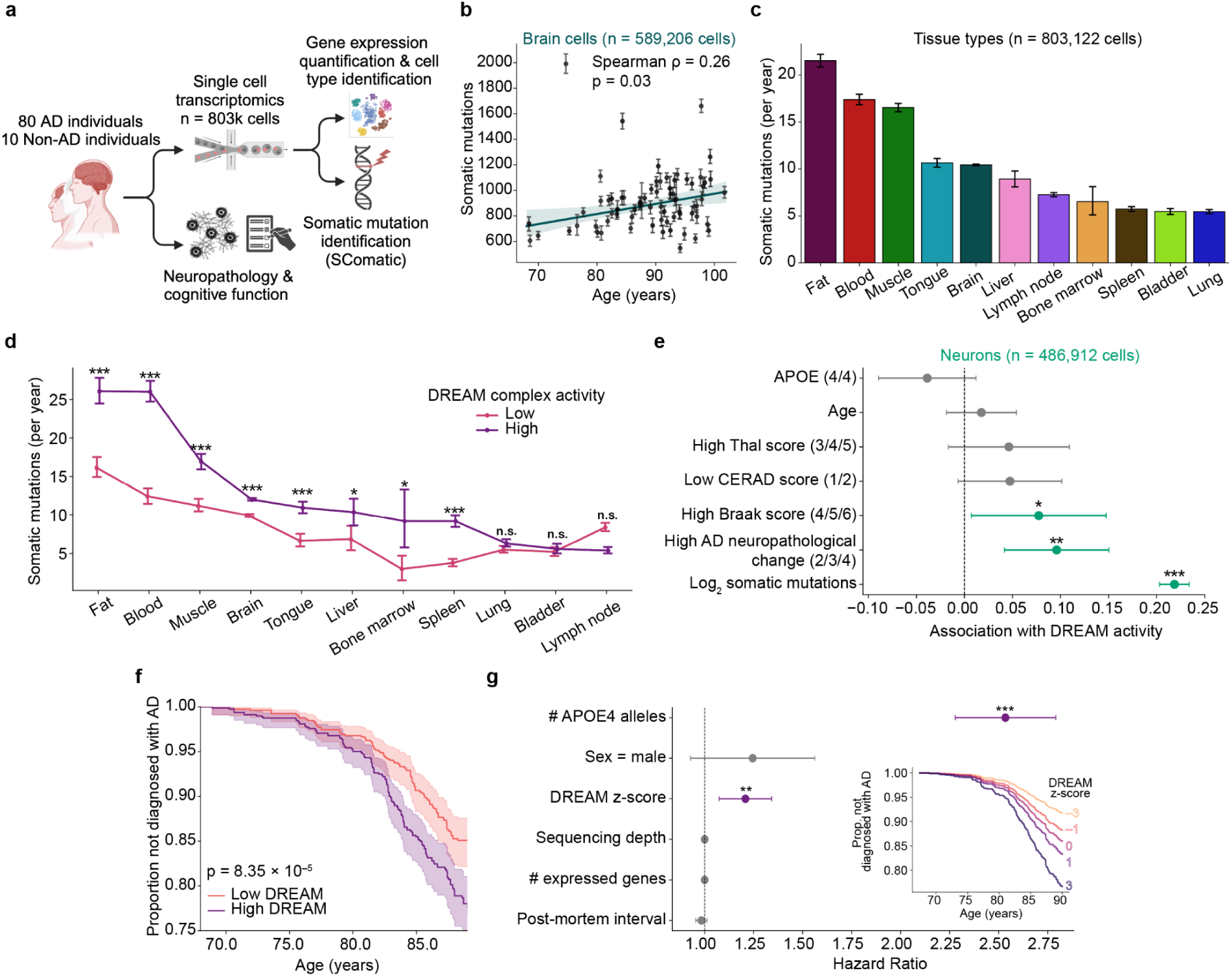
Multiple associations between DREAM activity and Alzheimer’s Disease. **a)** 90 human individuals (80 with AD or neurodegeneration and 10 cognitively normal) were profiled for mRNA expression in single cells from 11 tissues (n = 803,122 total cells, 73% of cells were from the brain) as well as neuropathology and cognitive function. Cell type and somatic mutation profile of each cell were identified from the single-cell mRNA. **b)** Scatter plot of the number of somatic mutations per haploid genome (**Methods**) identified in the brain of each individual, compared to the age of that individual (n = 77 individuals with ≥ 5,000 sequenced brain cells, n = 589,206 cells). Points and error bars indicate the mean and 95% confidence interval across all cells from an individual. P value calculated by modeling Spearman ρ’s as a Student’s t distribution. **c)** Barplot indicating the mean and 95% confidence interval of the number of somatic mutations identified in each cell per year within each tissue type (normalized to haploid genome size, n = 803,122 cells, n = 90 individuals). **d)** Number of somatic mutations in each cell (normalized to haploid genome size, n = 803,122 cells, n = 90 individuals, **Methods**) stratified by DREAM complex activity (highest vs. lowest 20% of DREAM activity scores within each tissue). Error bars denote 95% confidence intervals. (***), (*), and (n.s.) indicate p values from a one-sided Mann-Whitney test of p < 5.0 × 10^-17^, p < 0.005 (Bonferroni corrected p-value), and p ≥ 0.005, respectively. **e)** Association of cell or individual-level covariates with DREAM complex activity in neurons (n = 486,912 neurons, n = 77 individuals). Dots and whiskers indicate the coefficient and 95% confidence interval from linear mixed effects modelling (**Methods**). (***), (**), and (*) indicate p values from a two-sided t-test of p < 5.0 × 10^-30^, p < 5.0 × 10^-4^, and p < 0.05, respectively. Covariates with insignificant associations are shown in grey. **f)** Kaplan-Meier survival curve comparing the age at AD diagnosis between individuals with the top and bottom third of DREAM complex activities (purple vs. pink, n = 1,101 individuals). The dark line and shaded area indicate the proportion of individuals without AD at a particular age and the 95% confidence interval of this estimate, respectively. P value calculated using a two-sided log-rank test. **g)** Dot-and-whisker plot indicating the association of each variable on the y-axis with age at AD diagnosis (n = 1,101 individuals). Dots and whiskers indicate the hazard ratio and 95% confidence intervals from a Cox proportional hazard model (**Methods**). Asterisks indicate p values from a two-sided Wald test; (***), p < 5.0 × 10^-10^ and (**), p < 0.01. Covariates with insignificant associations are shown in grey. Inset: Line plot of the predicted effect of different levels of DREAM on age at AD diagnosis, holding other factors constant.

Neuropathological data had been collected at the time of death for Alzheimer’s diseased individuals (n = 80). We found that increased Braak score (a measure of tau pathology), Alzheimer’s Disease Neuropathologic Change score (ADNC, a composite measure of amyloid plaques, neurofibrillary tangles, and neuritic plaques), and somatic mutation burden were all associated with greater DREAM activity (**Fig. 5e, Methods**). In an independent dataset of post-mortem brains from individuals with neurodegeneration (Religious Orders Study/Memory and Aging Project, ROSMAP^44^, n = 1,101 individuals), elevated DREAM activity was associated with earlier AD diagnosis (**Fig. 5f**). Controlling for covariates, a one standard-deviation increase in DREAM activity corresponded to a 21% increase in the risk of AD at a given age (HR = 1.21, p = 0.01, **Fig. 5g**). Unlike DREAM-target genes, the expression of randomly selected sets of genes did not exhibit a significant association with AD diagnosis risk (mean HR = 1.0, **Supplementary Fig. 6h-i**). We also considered that reverse causation was possible: the longer an individual had AD before death, the lower their DREAM activity would become. However, DREAM activity was not associated with the duration of AD before death **(Supplementary Fig. 6j**).

These results indicated that DREAM activity is associated with human dysfunction at multiple biological scales: somatic mutations at the genomic level, neuropathology at the cellular level, and Alzheimer’s disease at the organismal level.

### DREAM loss of function reduces mutation accumulation *in vivo*

Finally, we tested whether DREAM activity is a causal driver of mutation accumulation *in vivo* by preventing DREAM complex assembly in mice and quantifying lifetime somatic mutation burden. We analyzed formalin-fixed (FFPE) brains from DREAM loss-of-function (LoF)^45^ mice (n = 4, p107^D/D^ p130^−/–^) and littermate controls (n = 5; p107^D/D^ p130^fl/fl^, **Fig. 6a**). At 8 weeks of age, both groups had received tamoxifen, preventing DREAM assembly only in the DREAM LoF mice, while littermate controls retained intact DREAM function^45^. Brain tissue was collected after natural death and profiled for somatic mutations using single-molecule duplex sequencing (UDSeq, **Methods**).

**Figure 6:**
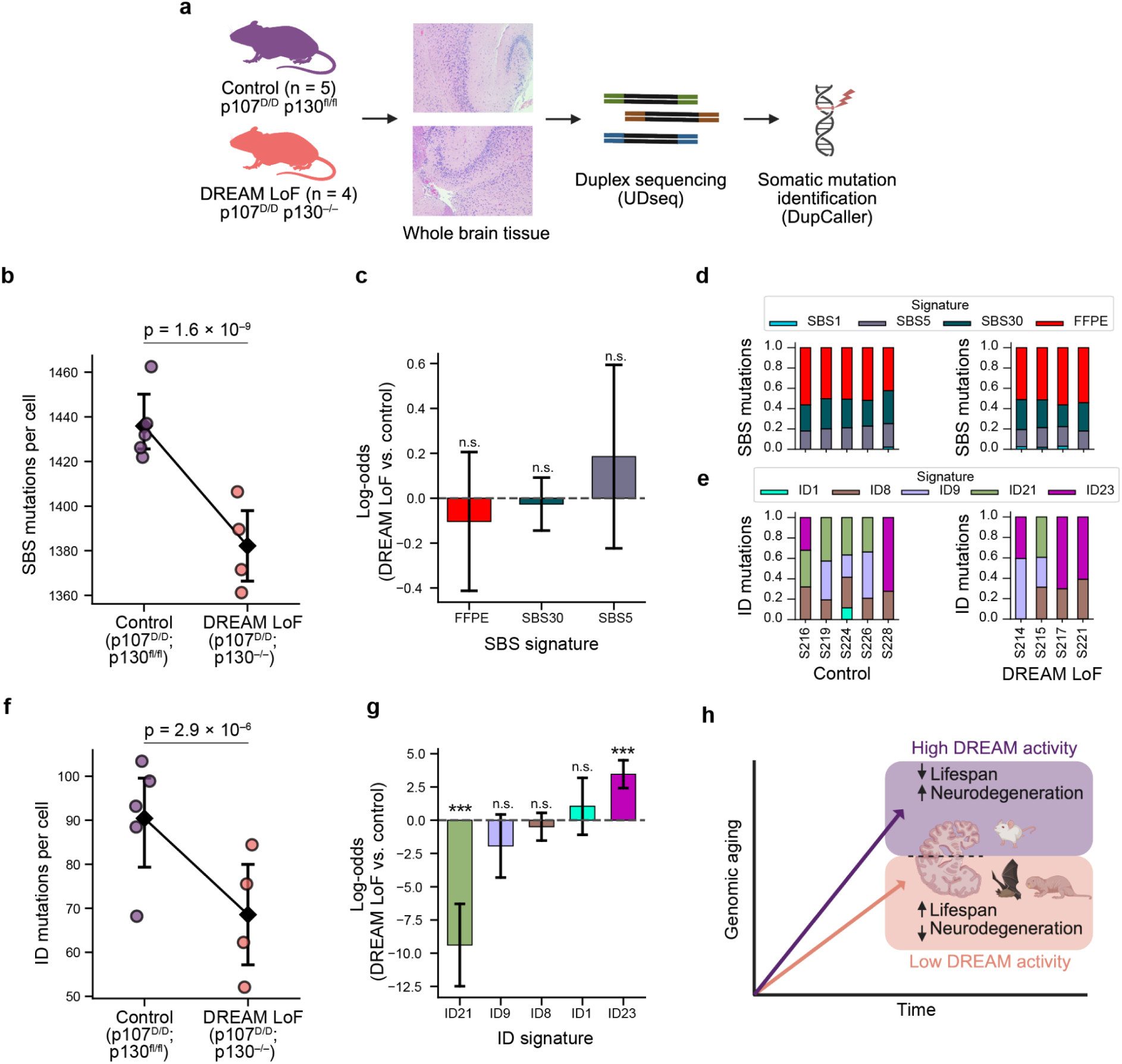
Reduced mutagenesis in brains of DREAM loss-of-function mice. **a)** Whole brain tissue slices were collected from control mice (p107^D/D^ p130^fl/fl^, n = 5) or DREAM loss-of-function mice (LoF, p107^D/D^ p130^−/–^, n = 4). Somatic mutations were identified via duplex sequencing (UDseq) and somatic variant calling (DupCaller). **b)** Single base substitution (SBS) mutations per (diploid) cell in each mouse. Points are individuals; diamonds and whiskers denote the means and 95% confidence intervals (CI). P value reflects the association between DREAM status and the count of SBS mutations per cell according to a negative binomial regression model (**Methods**). **c)** Log-odds of each SBS signature in DREAM LoF vs. control mice. Error bars indicate 95% CI. P values indicate significance of association between DREAM status and the proportion of mutations assigned to each signature from a two-sided binomial model (**Methods**), with n.s. corresponding to p > 0.05. **d-e)** Proportional contributions of (d) SBS signatures and (e) ID signatures (colored stacked bars) in each control or DREAM LoF mouse (x axis). The tissue preservation process (FFPE, formalin-fixed paraffin-embedded) is associated with a distinct SBS mutational signature. **f)** Similar to (b), but showing insertion/deletion (ID) mutations. **g)** Similar to (c), but for insertion/deletion (ID) mutational signatures. (***) and (n.s.) indicate p < 5.0 × 10^−9^ and p > 0.05, respectively. **h)** Model for the relationship between DREAM activity and aging phenotypes, in which higher DREAM activity associates with a faster rate of genomic aging due to repression of DNA repair. Consistent with this model, short lived species have higher DREAM activity than long lived species, and individuals with higher DREAM complex activity have accelerated neurodegeneration.

Controlling for covariates, we found that DREAM LoF mice exhibited a 4.2% reduction in single-base substitution (SBS) mutations per cell compared to controls (**Fig. 6b**, **Methods**). Across samples, the operative mutational signatures – distinctive mutation patterns that reflect the underlying source of somatic mutations^46^ – were SBS1, SBS5, SBS30, as well as the characteristic mutational signature of formalin fixation^47^. SBS1 reflects spontaneous deamination of 5-methylcytosine and SBS5 is a ubiquitous clock-like process of uncertain origin – both are classical age-related signatures – whereas SBS30 is linked to base-excision repair defects^48^. The proportion of mutations attributed to each mutational signature was not significantly different between groups, suggesting that loss of DREAM reduces overall mutagenesis rather than selectively altering a single repair pathway (**Fig. 6c-d**).

We observed that DREAM loss-of-function also conferred protection from insertion and deletion mutations (ID). Relative to controls, DREAM LoF mice had 19.6% fewer ID mutations (**Fig. 6e-f**). The ID mutational spectra also shifted: the prevalence of the ID21 mutational signature – 2-4 bp deletions in double-repeats – was significantly reduced in DREAM LoF brains, whereas ID23 – 1 bp deletions at homopolymer runs – was increased (**Fig. 6g**). Together, these results indicated that loss of DREAM activity reduces the lifetime accrual of somatic mutations, spanning multiple mutation types and mutational processes.

## Discussion

Collectively, our results suggest that the DREAM complex plays a pivotal role in modulating the rate of DNA damage accumulation, thereby shaping organismal lifespan and the onset of aging-related pathologies (**Fig. 6h**). Consistent with a causal role, genetic disruption of DREAM assembly protects mice from somatic mutation, directly linking DREAM activity to mutation accumulation in vivo (**Fig. 6**). These results corroborate, and may help explain, previous research demonstrating that long-lived species and centenarians possess mechanisms enhancing genomic stability^6,10–12^.

The simplest interpretation of our findings is that when the DREAM complex represses DNA repair, DNA damage tends to remain unresolved leading to somatic mutation and other consequences. Some of our experimental data indeed support this model: for instance, loss of DREAM reduced both SBS and ID mutation rates and reshaped the ID mutational spectrum (**Fig. 6**, reducing signature ID21^49^) consistent with enhanced repair of replication-slippage intermediates. Furthermore, DREAM target genes include those involved in nearly every major DNA repair pathway^23,24^, including MUTYH, NEIL3, and POLD1 (base excision repair^50,51^); RAD51, XRCC2, and BRCA2 (homologous recombination^52^); XRCC4, PARG, and MRE11 (non-homologous end joining^53^); and POLQ, PCNA, and USP1 (translesion synthesis^54^). Simultaneously, unrepaired lesions can give rise to mutations by multiple mechanisms – such as cytosine deamination in the case of UV-induced cyclobutane pyrimidine dimers^55^, mispairing of oxidized bases during DNA replication^56,57^, and error-prone translesion synthesis at replication forks stalled by DNA damage^58^. Taken together, these observations suggest that by reducing the expression of DNA repair factors, the DREAM complex allows DNA damage to persist, thereby increasing the likelihood that unrepaired lesions will be converted into permanent mutations through replication or error-prone repair processes.

Long-lived species exhibit a range of adaptations thought to underlie their extended lifespans, including enhanced DNA repair capacities^2^, reduced somatic mutation rates^4,5^, altered metabolic rates^59^, and larger body sizes^60^. We observed an inverse association between DREAM complex activity and species lifespan (**Fig. 3**), which may be either a cause or result of such adaptations. By permitting increased expression of DNA repair genes, reduced DREAM activity in long-lived species could plausibly enhance DNA repair and decrease somatic mutation rates. This idea aligns with reports of elevated expression of DNA repair genes in long-lived species^61,62^, although structural modifications to the repair factors themselves have also been implicated in such heightened repair capabilities^6^. Alternatively, reduced DREAM activity may be a downstream effect of other adaptations that more directly confer extended longevity. Although the observed relationship between DREAM activity and lifespan does not appear to be confounded by species size (**Fig. 3d**, **Supplementary Fig. 3a-b**), the possibility of other confounding factors, such as lower metabolic rate – or additional, yet unidentified mechanisms – warrants further investigation.

Within species, we found that elevated DREAM activity was associated with both the decreased survival of mice fed a high-fat diet (**Fig. 4**) and earlier onset of AD in humans (**Fig. 5**), a pattern potentially explained by heightened genomic damage. Indeed, high DREAM activity correlated with greater somatic mutation burdens and more severe neuropathology, aligning with prior evidence implicating DNA damage in age-related diseases^63–65^ in general and in Alzheimer’s disease^66,67^ in particular. Nevertheless, we cannot exclude the possibility that in these cases DREAM activity is merely a marker of other processes that drive morbidity and mortality, rather than a direct cause.

Extrapolating from these findings, DREAM inhibition could represent a new therapeutic approach to prevent or mitigate DNA damage-driven conditions, including aging. Compounds which disrupt DREAM complex formation^23,68,69^ have already shown beneficial effects in model organisms – reducing oxidative stress^70^, preserving bone density^71^, enhancing insulin sensitivity^72,73^, and improving cognitive function^74^. Our results suggest that the upregulation of DNA repair brought about by these DREAM inhibiting compounds may underlie their therapeutic benefits. While some of these compounds are in early development for human neurological disorders^75,76^, the link between DREAM activity and DNA repair explored here points to a broader opportunity to use DREAM-modulating therapies to slow or prevent age-related pathologies in humans.

## Methods

### scRNA-seq processing

Each scRNA-seq dataset was processed separately using Scanpy^77^. First, any cell with fewer than 200 detected genes or more than 10,000,000 total read counts was removed. Second, any gene detected in fewer than three cells was discarded (using the Scanpy functions ‘filter_cells’ and ‘filter_genes’). Third, the total read count within each cell was normalized to 10,000 (‘normalize_per_cell’), log-scaled (‘log1p’), and then scaled to have a value between zero and ten (‘scalè).

### Calculation of DREAM activity

DREAM-regulated genes (n = 328 genes) were defined as genes with promoters concurrently bound by at least 3 core DREAM complex subunits in a ChIP-seq experiment targeting LIN9, LIN54, p130, and E2F4 in quiescent human cells^24^. The Single Sample Gene-Set Enrichment (ssGSEA^78^) of the expression of these genes was calculated in each RNA-seq or scRNA-seq sample. For datasets in which the expression of one or more DREAM regulated genes was not detected, the remaining genes were used in the ssGSEA calculation. The ssGSEA “normalized enrichment score” was regressed against the number of expressed genes and total count of reads in each cell/sample, to correct for technical variation arising from sequencing depth^79^. The residuals from this regression (i.e., the normalized enrichment score corrected for sequencing depth) were inverted and scaled to be positive, making higher scores correspond to greater DREAM activity (i.e., lower expression of DREAM-regulated genes). The resulting value was termed “DREAM activity” (**Fig. 1**).

### Proteomic DREAM activity validation

Matched transcriptomic and proteomic data from the Clinical Proteomic Tumor Analysis Consortium (CPTAC)^80^ were used to compare the transcriptomic DREAM activity score with the abundance and phosphorylation state of LIN52 and p107 (RBL1). A total of 160 individuals were included from the clear cell renal cell carcinoma (ccRCC) and head and neck squamous cell carcinoma (HNSCC) cohorts; samples in which the DREAM components were not detected were excluded. Multivariate linear regression was performed using the model:

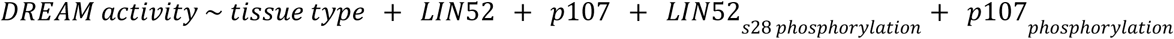

where *p*107_*phosphorylation*_ was the mean phosphorylation across 17 detected phospho-sites on p107.

### Identification of somatic mutations in scRNA-seq

For the Tabula Sapiens^42^ and SEA-AD^81^ sc/snRNA-seq datasets, SComatic^41^ was applied with default parameters to identify somatic single nucleotide substitution mutations. In brief, first somatic mutations in each “pseudo-bulked” cell type in each individual were identified, treating variants present in more than one cell type as germline. Second, somatic mutations in each individual cell were identified and filtered to include only variants also identified at the “pseudo-bulk” level of the respective cell type. Third, the count of somatic mutations in each cell was divided by the number of basepairs from which mutations were called in that cell – due to there being sufficient coverage at these loci – to define the somatic mutation rate per base pair. Cells without sufficient coverage for mutation identification in greater than 10 kb were discarded. The approximate number of somatic mutations per cell was then calculated by multiplying the somatic mutation rate per base pair by the approximate number of base pairs in the haploid human genome (3.0 × 10^9^). Finally, the somatic mutation rate per cell per year was calculated by dividing the number of somatic mutations per cell by the age of the donor of that cell. For the Tabula Muris Senis dataset, somatic mutations had been identified by the original publication^33^, and we simply scaled these somatic mutation burdens to be relative to the number of bases from which mutations were called.

### Somatic mutation rate validation

The somatic mutation rate per kb per month, estimated from scRNA-seq in mouse skin cells (**Fig. 2**), was compared with a previous estimate of somatic mutation rates in mouse fibroblasts^7^ obtained using the single-cell multiple displacement amplification (SCMDA) DNA-sequencing method^82^. Milholland et al.^7^ found there to be approximately 4.4 × 10^-7^ somatic mutations per bp in a 5 day old mouse. We converted this estimate to be in units of mutations/kb/month, using the average month length of 30.44 days, and found our estimate of 0.0030 mutations/kb/month to be remarkably similar to this previous estimate of 0.0027 mutations/kb/month.

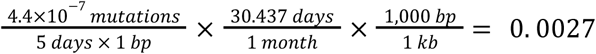

We also compared the mutation rates we identified from human sc/snRNA-sequencing (**Fig. 5**) to previous estimates based on DNA sequencing of the same or similar cell types as those considered in our analyses. This included seven estimates of mutation rates in immune cells^83,84^, three in neurons^83,85^, and one in each of muscle satellite cells, skin fibroblasts, and cells from the bladder urothelium^83^. DNA- and RNA-based mutation estimates were generally consistent though the rates we now observed were marginally lower (**Supplementary Fig. 6b-f**). This difference in mutation rates may reflect that RNA, unlike previous measures from DNA, is primarily derived from coding sequences which have reduced somatic mutation rates relative to non-coding sequences due to increased mismatch and transcription-coupled repair^8,86–88^.

### Generation of random background distributions

To serve as background distributions for the association of DREAM activity with somatic mutation rate (**Fig. 2g-h**), species maximum lifespan (**Fig. 4c**), and age at AD onset (**Supplementary Fig. 6h-i**), the association between the expression of random gene sets and these measures was calculated. In each dataset, 500 – 1,000 sets of genes were randomly selected, with replacement, from all genes detected in that dataset. The number of genes in each of these random sets was defined to be the same as the number of DREAM regulated genes detected in that dataset (n = 249 for the TMS mouse scRNA-seq, n = 308 for the cross-species RNA-seq dataset, and n = 322 for the ROSMAP human brain RNA-seq). The “activity” of these random gene sets was then calculated by treating these genes as if they were DREAM regulated genes and applying the process that was used to calculate DREAM activity (i.e., ssGSEA followed by sequencing depth correction and inversion, see the ‘Calculation of DREAM activity’ methods section). Finally, the activity of these random gene sets was compared to the respective measures of interest and the strength of that association was contrasted to that of DREAM activity and the measure of interest.

### Comparison of lifespan across species

Cross-species comparisons of DREAM activity, maximum lifespan, adult weight, and somatic mutation rate were conducted using a dataset of gene expression data (RNA-seq) collected from 3 tissues (brain, liver, kidney) of 92 different species^35^ (n = ∼3 samples per species per tissue, n = 803 total samples). We first calculated a representative DREAM activity for each species in each tissue by averaging the DREAM activity scores of all individuals of a given species in a tissue. Species with fewer than two sampled individuals in a given tissue were discarded from comparisons considering that tissue; taxonomic orders with only a single sampled species were discarded. We examined the association of DREAM activity with maximum lifespan (ML) using zero-intercept reduced major axis regression (RMA, implemented by the pylr2 regress2 function^89^), in each tissue separately (**Fig. 4b, c**). Because body size can confound interspecies lifespan relationships^90^, we conducted the same DREAM vs. ML comparison but correcting for adult body weight (AW). To do so, we performed an allometric regression of each trait (DREAM activity and ML) on log_10_-transformed AW and compared the resulting residuals (i.e., partial regression of DREAM activity vs. ML with respect to AW, **Fig. 5d, Supplementary Fig. 3a-b**). Finally, integrating a previously published estimate of somatic mutation rates^4^ for a subset of the same species, we performed analogous cross-species comparisons of DREAM complex activity and somatic mutation rate, both with and without adjusting for AW (**Fig. 4f**, **Supplementary Fig. 3c-g**).

### Survival analysis of mouse strains

The Williams et al.^40^ dataset consists of 50 distinct strains of inbred mice split into two cohorts: the “Deeply Profiled Cohort,” which was profiled for liver gene expression (RNA-seq), blood biomarkers, and the weight of different organs (n = 270 mice); and the “Lifespan Cohort,” which was followed to measure the lifespan of each strain (n = 612 mice). The DREAM activity of each mouse in the Deeply Profiled Cohort was calculated from the liver RNA-seq. Four strains with highly variable DREAM activity scores across mice were removed (standard deviation > 0.2). The Deeply Profiled Cohort was used to estimate strain-specific covariates for use in a Cox proportional hazards model (lifelines CoxPHFitter^91^). This model was fit to the survival data of the Lifespan Cohort to measure the association of lifespan with diet (high fat vs. chow diet), DREAM activity, weight, clinical biomarkers, and weight change (**Fig. 4**).

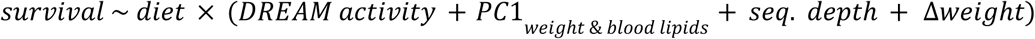

Covariates in the Cox model included: the mean DREAM activity value within each strain on each diet; the mean sequencing depth in each strain, to account for any confounding of DREAM activity by differences in sequencing depth between strains; the first principal component (PC1, 25.2% of variance explained, **Supplementary Fig. 5g**) from an analysis of normalized clinical biomarkers and organ weights (variables listed in **Supplementary Fig. 5g**); diet group (chow or high fat diet); and change in body weight since starting the respective diet.

### Neuropathology vs. DREAM activity

Neurodegenerative pathology was compared with DREAM complex activity using the SEA-AD^81^ dataset, in which snRNA-seq was carried out on 803,112 brain cells from 83 individuals. Neuropathological assessments, including Thal phase, CERAD score, and Braak stage, had been conducted for each individual and integrated into an overall AD neuropathological change score^92^ (ADNC). Each variable was binarized into high (Thal ≥ 3, Cerad ≤ 2, Braak ≥ 4, ADNC ≥ 2) versus low AD risk. Then, linear mixed effect regression was used to test the association between each of these variables – in addition to APOE 4/4 genotype, age (scaled to units of decades), and the count of somatic mutations per cell (log_2_ scaled) – and DREAM activity (**Fig. 5e**). These variables were treated as fixed effects, while the individual identifier was modeled as a random effect, to account for the nested structure of the data, with multiple cells sampled per individual.

### Association of DREAM activity with age at AD diagnosis

The ROSMAP^44^ (Religious Orders Study and Rush Memory and Aging Project) dataset was used to investigate the association of DREAM activity with the diagnosis of Alzheimer’s disease. In this study, RNA-sequencing data were collected from multiple brain regions (dorsolateral prefrontal cortex, head of the caudate nucleus, posterior cingulate cortex, temporal cortex, and frontal cortex) of deceased donors (mean 2.4 ± std 1.0 brain regions per individual, n = 1,105 individuals, n = 2,776 samples). A Cox proportional hazard model (lifelines CoxPHFitter^91^) was fit to predict the age at AD diagnosis of these individuals. Age at AD diagnosis was right-censored at 90 years to ensure masking of donor identity. The DREAM activity, number of APOE 4 alleles, sex, total RNA sequencing depth, count of genes with detected expression, and the duration of time between death and tissue procurement (post-mortem interval) were used as covariates in the model. DREAM activity scores were Z-score normalized within each brain region and outliers were dropped (samples with DREAM |Z| > 5). As there were multiple samples from each individual, the model was corrected for intra-individual correlations through clustered variance adjustment (**Fig. 5f-g**).

### DREAM deficient mice

We leveraged a previous study^45^ which had engineered mice with inactivated DREAM activity. In these mice, DREAM assembly was prevented by combining a constitutive *Rbl1* missense allele (p107^D/D^), which disrupts MuvB binding, with tamoxifen-induced UBC-Cre-ERT2 deletion of *Rbl2*. Because either p107 or p130 can scaffold DREAM, loss of p130 in the p107^D/D^ background abolishes assembly of the complex^45^. Control mice were Cre-negative. At 8 weeks of age, both groups were treated with tamoxifen – this activated the deletion of gene encoding for p130 (*Rbl2*) producing DREAM loss-of-function (LoF) mice (p107^D/D^ p130^−/−^, n = 5), while the control mice retained intact DREAM activity due to Cre-negativity (p107^D/D^ p130^fl/fl^, n = 5). Brain tissue was obtained from these mice at death, formalin-fixed, and paraffin-embedded.

### Duplex sequencing (UDSeq) library preparation and sequencing

Formalin-fixed paraffin-embedded (FFPE) mouse brain tissues (n = 10) were sectioned, stained with hematoxylin and eosin, and punches containing sufficient nuclei were selected by microscopic inspection. Genomic DNA was extracted using the QIAamp DNA FFPE Advanced Kit (Qiagen, cat. no. 56704) according to the manufacturer’s instructions and quantified by Qubit, with 500 ng per sample normalized to 5 ng/µl for library preparation. Genomic DNA was enzymatically fragmented using the UltraShear system (M7634L) for 25 minutes, targeting an average fragment size of approximately 350 base pairs. The fragmented DNA was then processed using the UDSeq library preparation workflow with the xGen™ cfDNA & FFPE DNA Library Preparation Kit. All steps were performed on magnetic beads to minimize DNA loss during purification and enhance library conversion efficiency. For final library construction, 0.175 femtomoles of UMI-ligated DNA were amplified via 15 cycles of PCR to incorporate Illumina®-compatible dual index sequences. For matched normal samples, 5 femtomoles of UMI-ligated DNA from the same source were amplified using 10 PCR cycles. The resulting universal dual-indexed libraries were sequenced at the UCSD IGM Genomics Center using a 25B flow cell on the NovaSeq X platform.

### Identification of somatic mutations in duplex sequencing data

Sequencing data were downloaded and analyzed within the Triton Shared Computing Cluster at the San Diego Supercomputer Center (https://doi.org/10.57873/T34W2R). Somatic mutation calling and mutational burden analysis were performed using DupCaller 1 (v1.0.1)^93^ on UDSeq data with matched normal samples. Variant Call Format (VCF) files generated by DupCaller were used for mutational profiling and signature assignment, following our previously established methodology using the SigProfiler suite of tools. In particular, SigProfilerAssignment was run on the variant call set, with the entire mouse COSMIC catalog and FFPE signature from Guo et al. 2022^47^ FFPE signature as the input signatures. Mutations classified as FFPE artifacts were discarded and remaining mutations were analyzed. For each sample, the raw mutation count in the sequenced bases was extrapolated to estimate the number of mutations expected if the entire diploid genome had been covered at sufficient depth, yielding a per-cell equivalent mutation burden (termed “mutations per cell”).

### Analysis of UD-seq mutation calls

Negative binomial regression models were used to test for a difference in frequency of single base substitutions and insertion/deletion mutations between DREAM LoF and control mice. Models of three complexities were compared:

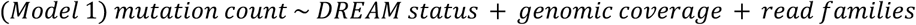

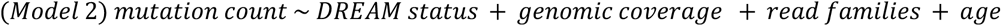

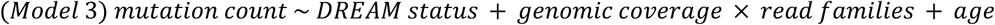

Model 3 showed the best fit across all mutation types as measured by log-likelihood (**Supplementary Fig. 7a-f**) and was therefore used for all statistical comparisons of mutation frequency. For comparisons of the proportional contribution of mutational processes, a binomial model of the same form was used. One outlier sample – identified by having a Cook’s distance^94^ > 1 and total mutation count |Z-score| > 3 across both mutation classes – was removed (**Supplementary Fig. 7g-i**).

### Software

All analyses were performed in Python 3.10. Data analysis was conducted using Pandas 1.5.3, SciPy 1.10.0, Pingouin 0.5.3, and Statsmodels 0.13.5. Data were visualized with Seaborn 0.12.1 and Matplotlib 3.7.1. No statistical method was used to predetermine sample size, the experiments were not randomized, and the investigators were not blinded to allocation during experiments and outcome assessment. Specific statistical approaches used are noted in the respective methods sections and figure captions.

## Data availability

Raw UD-sequencing data can be accessed on the Sequence Read Archive (SRA) at accession number [To be uploaded before publication]. Other data can be accessed through the respective publications (see **Supplementary Table 1**).

## Code availability

All custom algorithms and analysis code are available in the GitHub repository at https://github.com/zanekoch/dream_proj.

## Acknowledgements

This study was funded by the National Institutes of Health under awards U54 CA274502 and P41 GM103504. DNA sequencing was conducted at the IGM Genomics Center, University of California, San Diego, La Jolla, CA. This publication includes data generated at the UC San Diego IGM Genomics Center utilizing an Illumina NovaSeq 6000 that was purchased with funding from a National Institutes of Health SIG grant (#S10 OD026929). The Triton Shared Computing Cluster (TSCC) provided the computational resources used for DNA sequencing analyses.

## Author Contributions

ZK designed the study, carried out the primary data analyses, and wrote the manuscript. ABD, DHM, BS, and LA designed the study and wrote the manuscript. FD and PP designed the study and provided the mouse samples. KL and SN designed the study, performed the duplex sequencing experiment, and wrote the manuscript. TI designed the study and wrote the manuscript.

## Supplementary Figures and Tables

**Supplementary Table 1.**
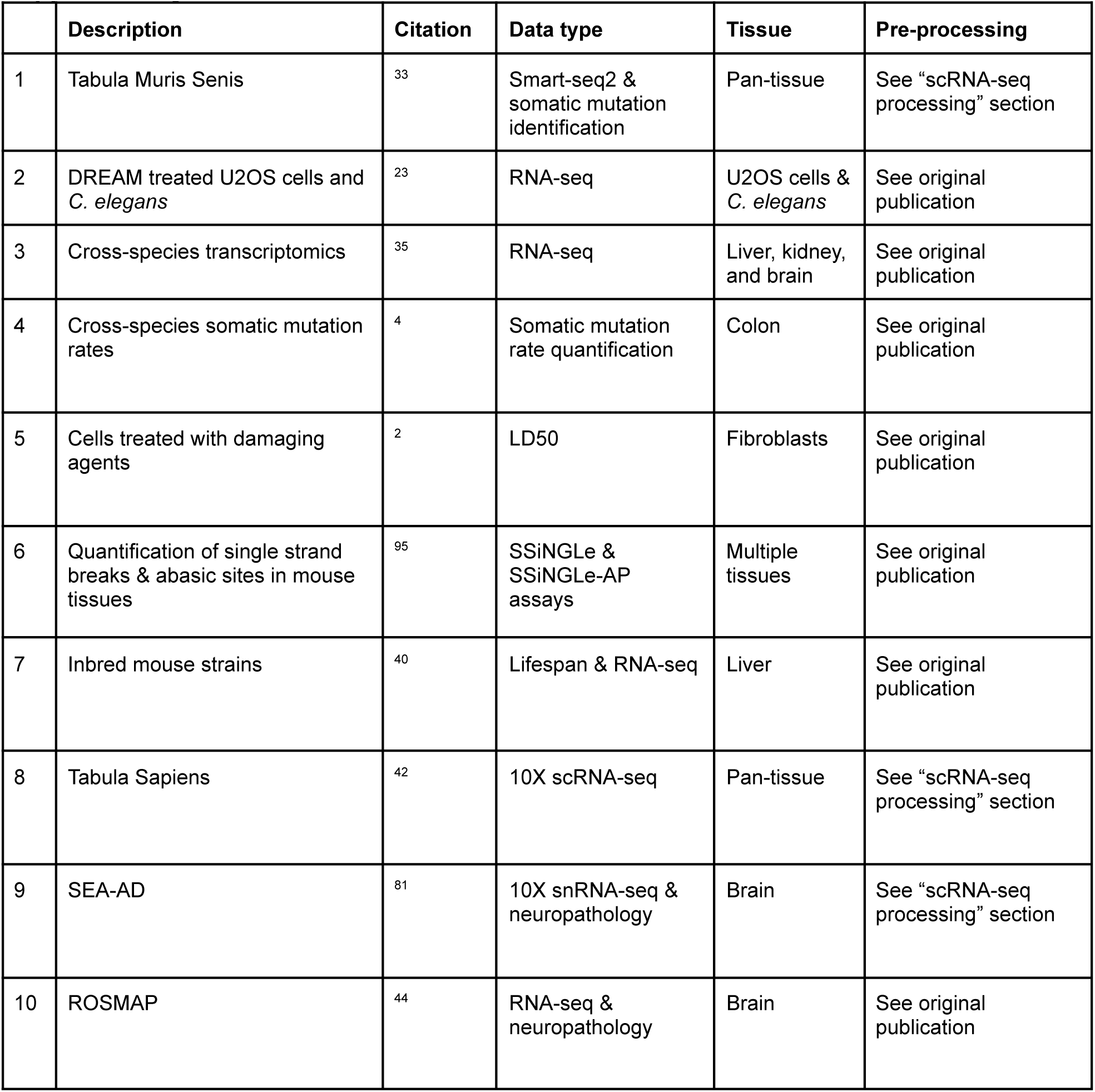

**Supplementary Figure 1:**
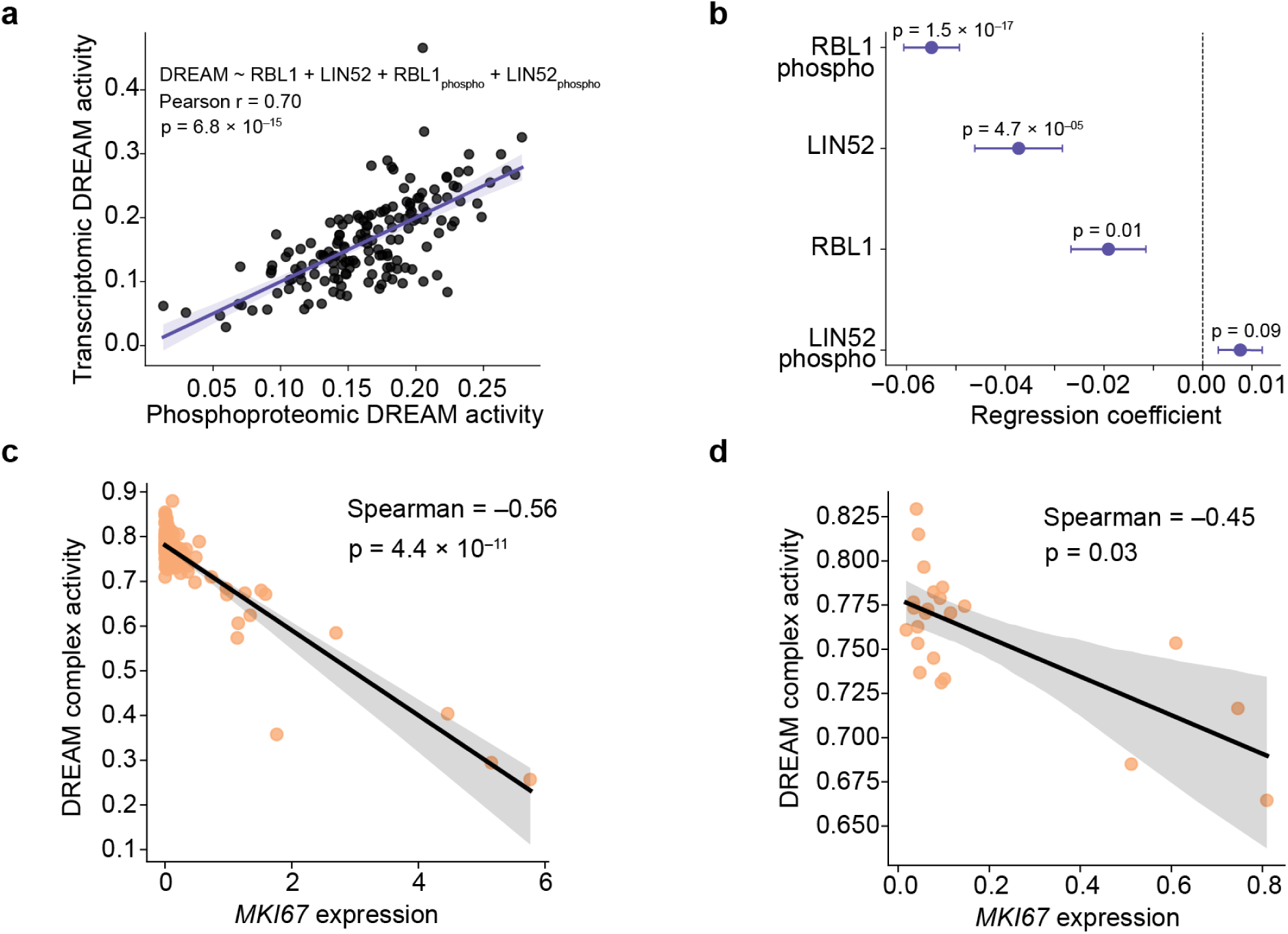
DREAM target expression mirrors other activity measures. **a)** Scatterplot of DREAM activity in human tumor tissues inferred from either the protein abundance and phosphorylation status of DREAM complex components (x-axis) or the gene expression of DREAM-target genes (“transcriptomic DREAM activity”, y-axis, n = 160 individuals). Two-sided p value calculated based on the exact distribution of Pearson’s r modeled as a beta function. **b)** Association of protein abundance and phosphorylation status of DREAM complex components (RBL1 and LIN52) with transcriptomic DREAM activity in human tissues (n = 160 individuals). Dots and whiskers indicate the coefficient and standard error from a linear mixed effects model regressing DREAM activity on the variables shown (**Methods**). Significance was calculated using two-sided t-tests. **c)** Scatter plot of the mean expression of the proliferation marking gene MKI67 versus the mean DREAM complex activity, in each cell type (n = 120 cell types). Two-sided p value calculated modeling Spearman ρ’s as a Student’s t distribution. **d)** Similar to (c) but in each tissue type (n = 21 tissues).

**Supplementary Figure 2:**
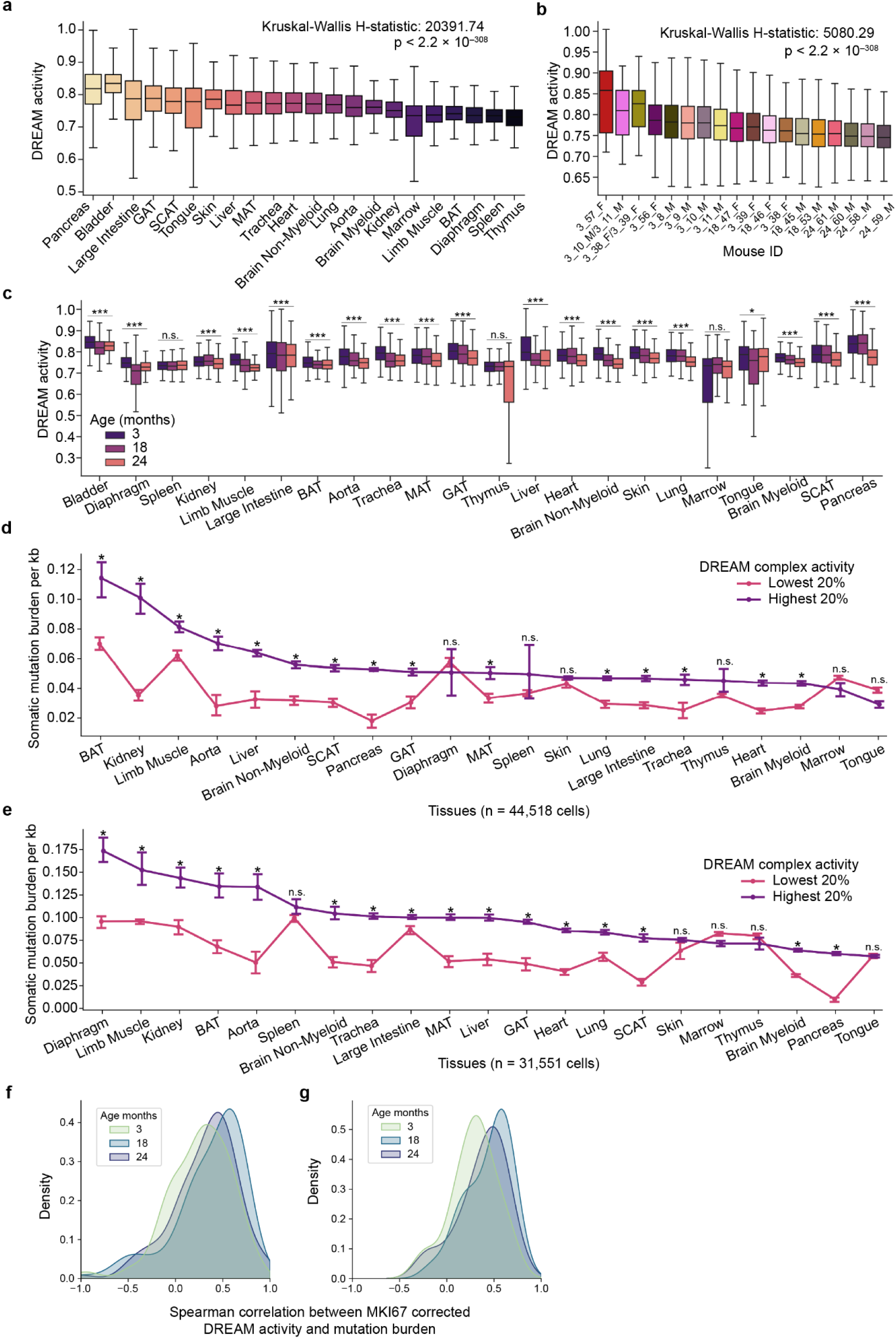
DREAM activity in single cells. **a)** Box plots of the distribution of DREAM complex activity by tissue across all cells from all mice (n = 110,824 cells, n = 18 mice). Two-sided p value from a Kruskal-Wallis test for a difference of distribution between tissues is shown. Boxes show inter-quartile range (IQR) with median line; whiskers extend to 1.5 IQR **b)** Box plots of the distribution of DREAM complex activity by mouse across all cells from all mice (n = 110,824 cells, n = 18 mice). Two-sided p value from a Kruskal-Wallis test for a difference of distribution between tissues is shown. **c)** Box plots of the distribution of DREAM activity scores in each tissue at each age (n = 110,824 cells, n = 18 mice). (***), (*), and (n.s.) indicate p values calculated by modeling Spearman ρ’s as a Student’s t distribution of p < 1.0 × 10^-7^, p < 0.01, and p ≥ 0.01, respectively. **d)** The somatic mutation burden per kilobase in cells (n = 44,518 cells, n = 10 mice) stratified by DREAM complex activity in 3 month old mice. Cells were partitioned into those with the highest and lowest 20% of DREAM complex activities within each tissue. Error bars denote 95% confidence intervals. (*) indicates p < 0.0023 (Bonferroni corrected p-value) and (n.s.) indicates p ≥ 0.0023, based on a two-sided Mann-Whitney test. **e)** Similar to (c), but for cells from 24 month old mice (n = 31,551 cells, n = 4 mice). **f)** The distribution of Spearman correlation coefficients between MKI67-corrected DREAM complex activity and somatic mutation burden, within each cell type and age (n = 120 cell types, n = 110,824 cells, **Methods**). **g)** Similar to (e) but within each tissue type (n = 21 tissues, n = 110,824 cells).

**Supplementary Figure 3:**
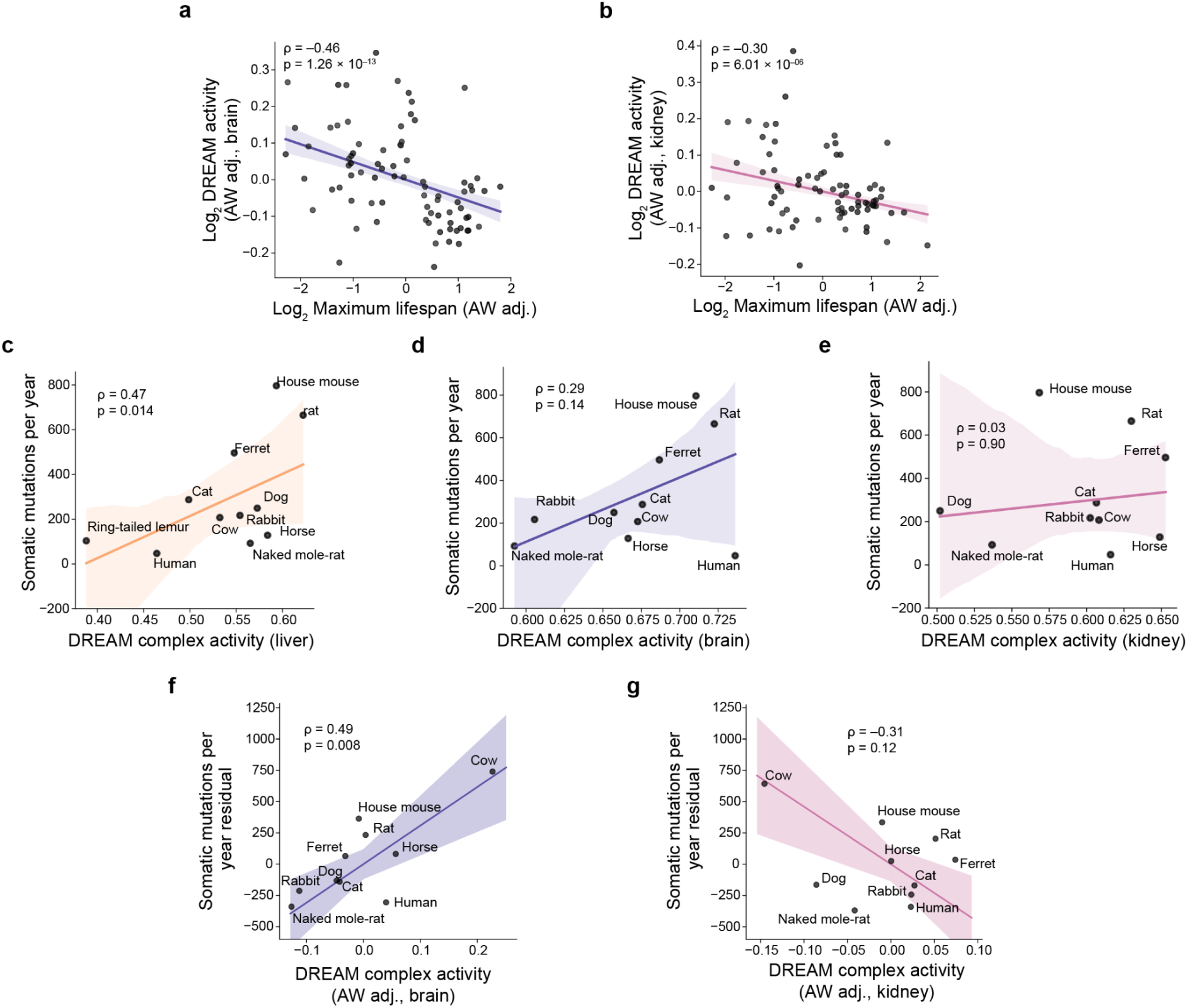
Mutation rate across species. **a)** Scatterplot and reduced major axis regression depicting the average adult weight (AW) adjusted DREAM activity and lifespan across species (n = 92 species, **Methods**). Both axes are shown on a logarithmic scale. **b)** Similar to (a), but DREAM activity is measured in the kidney. **c)** Scatterplot and reduced major axis regression of the mean DREAM activity in the liver of a species and somatic mutation rate in the colon in that same species (n = 11 species, **Methods**). **d)** Similar to (c), but DREAM activity is measured in the brain. **e)** Similar to (c), but DREAM activity is measured in the kidney. **f)** Scatterplot and reduced major axis regression depicting the average AW adjusted DREAM activity and somatic mutation rate (in the brain and colon, respectively) across species (n = 11 species, **Methods**). **g)** Similar to (f), but DREAM activity measured in the kidney.

**Supplementary Figure 4:**
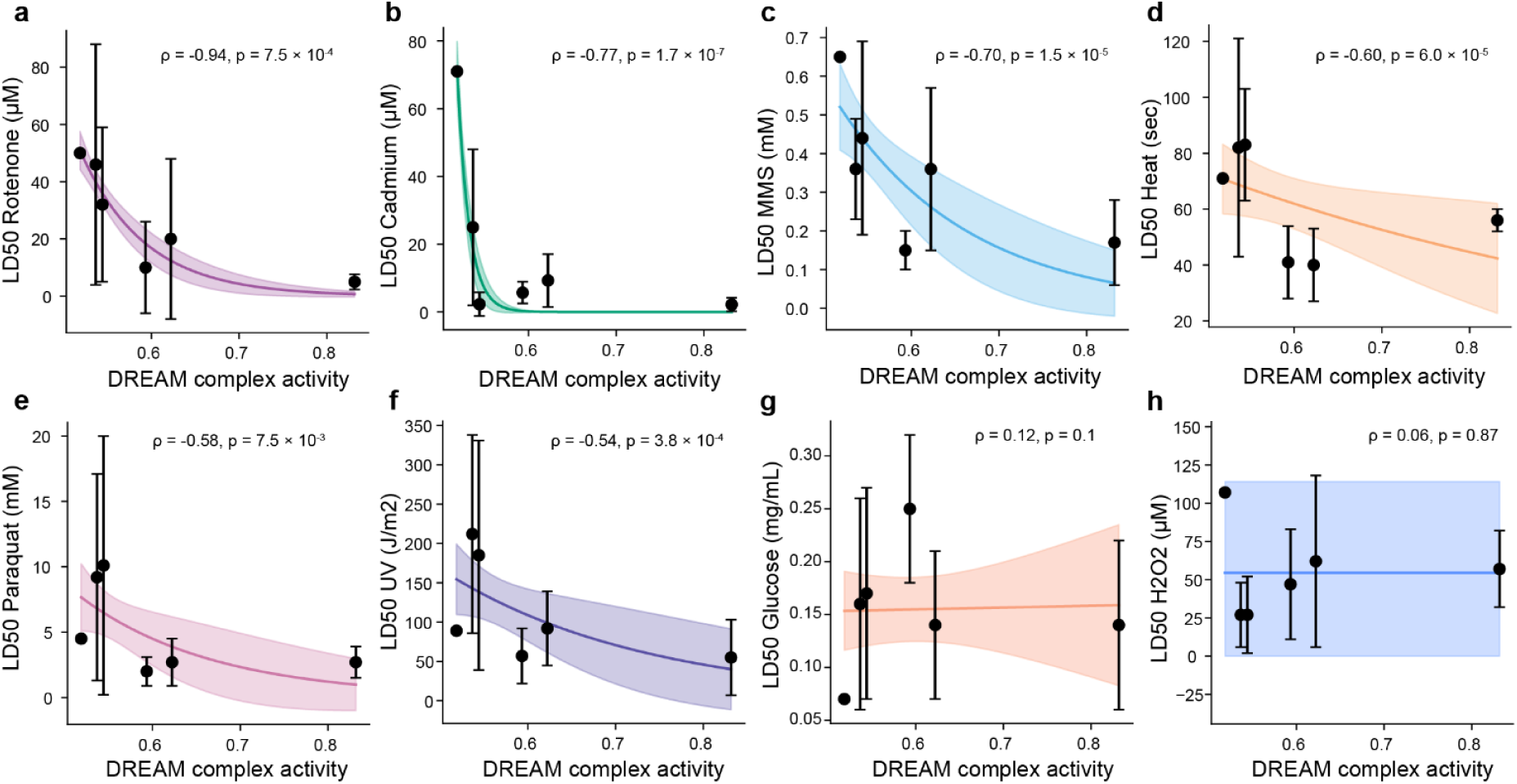
Connection of DREAM activity with DNA damage. **a-h)** Scatter plots of species-specific liver DREAM complex activity vs. resistance (LD50) of fibroblasts from six species to damaging agents (from Harper et al.^2^). The points and error bars indicate the mean and 95% confidence interval of the LD50 of all cell lines of a species. The colored line and shaded area indicate the fit of an exponential decay model and 95% confidence interval, respectively. From left to right in each plot species include: the north american beaver (n = 1 cell line), red squirrel (n = 9 cell lines), white-footed mouse (n = 7 cell lines), house mouse (n = 9 cell lines), Norway rat (n = 6 cell lines), and deer mouse (n = 5 cell lines). Significance was calculated using a two-sided t-test between the cell lines of the 3 species with highest and lowest DREAM activity, respectively.

**Supplementary Figure 5:**
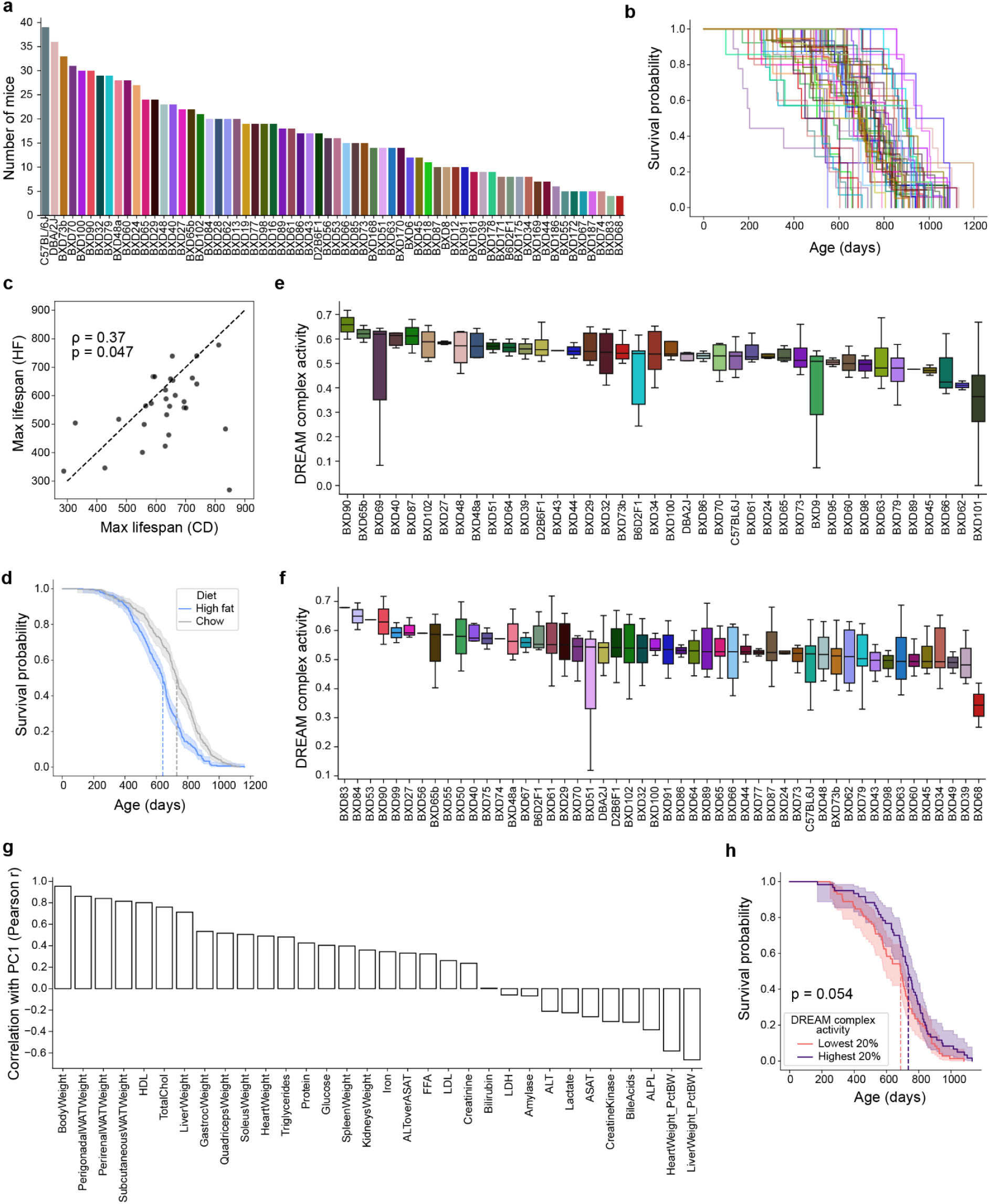
Survival and DREAM activity in mice. **a)** Bar chart of the number of mice of each strain (n = 50 strains, n = 882 mice). **b)** Kaplan-Meier survival curve for each strain in (a). **c)** Scatterplot comparing the maximum lifespan of each strain on the chow diet (CD, x-axis) to the high-fat diet (HFD, y-axis, n = 29 strains profiled for lifespan and RNA-seq on both diets). **d)** Kaplan-Meier survival curve for mice of all strains on the CD (n = 498 mice) compared to the HFD (n = 522 mice). The dark line and shaded area indicate the proportion of mice surviving to a particular age and the 95% confidence interval of this survival estimate, respectively. Dashed lines indicate the median survival of each group. **e)** Box plots of the distribution of DREAM complex activity in mice of each strain fed the HFD (n = 120 mice profiled for liver gene expression). **f)** Similar to (e) but for mice fed the CD (n = 150 mice profiled for liver gene expression). **g)** Correlation of the standardized values of each covariate with principal component 1 (i.e., “PC1 loading”, n = 270 mice, **Methods**). **h)** Kaplan-Meier survival curve of CD fed mice belonging to strains with the highest vs lowest 20% of DREAM complex activities (purple vs. pink, n = 132 mice, n = 15 strains). The dark line and shaded area indicates the proportion of mice surviving to a particular age and the 95% confidence interval of this survival estimate, respectively. Dashed lines indicate the median survival of each group. P value calculated from a two-sided log-rank test.

**Supplementary Figure 6:**
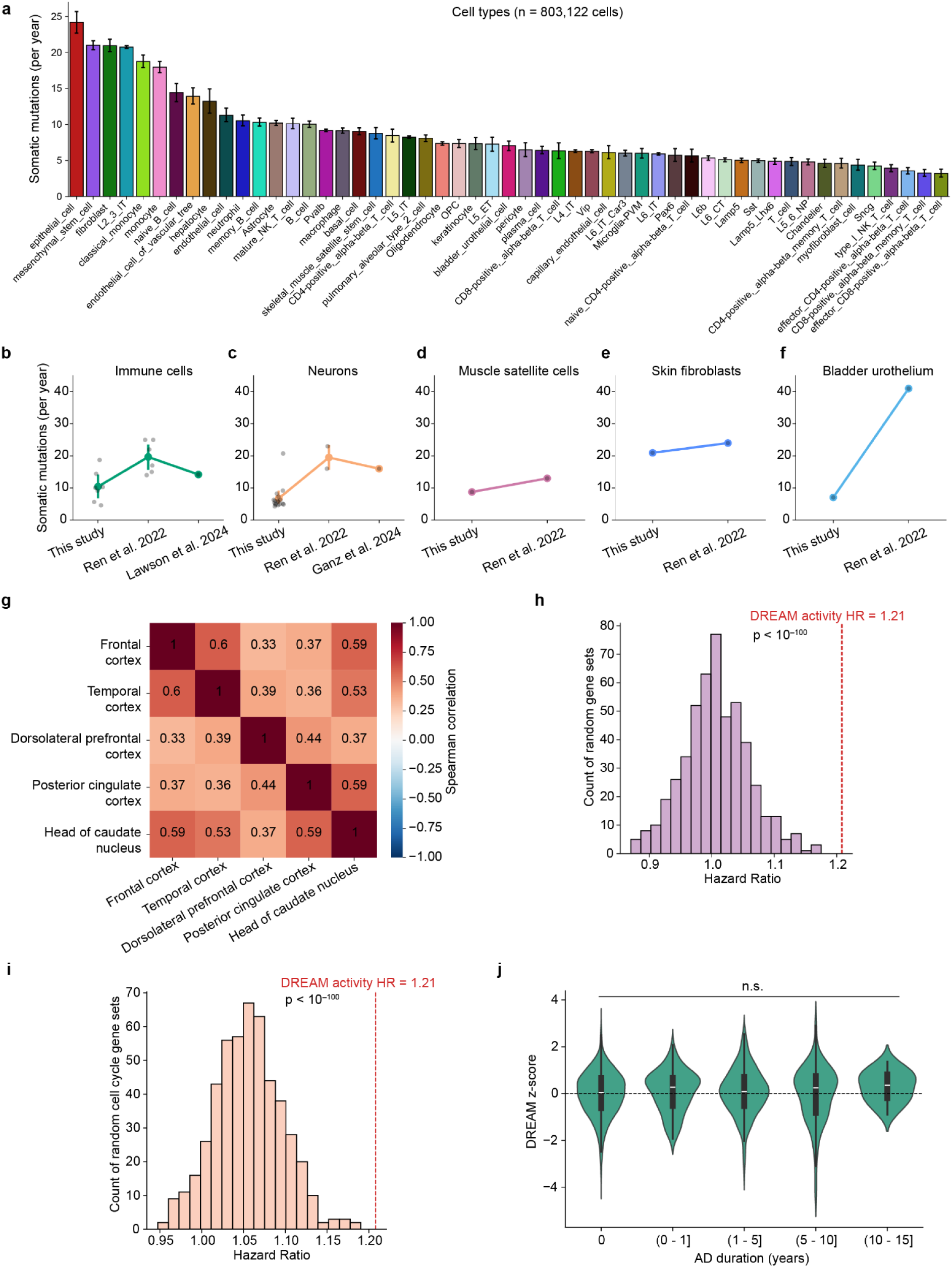
Somatic mutations in human tissues. **a)** Barplot indicating the mean and 95% confidence interval of the number of somatic mutations identified in each cell per year (normalized to haploid genome size, **Methods**) grouped by cell type (n = 803,122 cells, n = 90 individuals). **b-f)** Line plots comparing the number of somatic mutations per year identified by “this study” compared with previous estimates using DNA sequencing. Dark points indicate the mean mutation rate for a single cell type in a particular study, while the colored points indicate the mean and 95% confidence interval of these mutation rates. The x-axis lists the study from which the respective data came. **g)** Heatmap depicting the pairwise Spearman correlations of DREAM complex activity in different brain regions of the same individual (n = 1,101 individuals, 2,776 samples, mean 2.4 ± std 1.0 brain regions per individual). All pairwise correlations are significant (p < 0.0004). **h)** Histogram showing the distribution of hazard ratios from Cox proportional hazards models predicting age at AD diagnosis (using the same covariates as **Fig. 6g, n** = 1,101 individuals, **Methods**) using the ssGSEA enrichment of 500 randomly chosen sets of genes. HR of DREAM-regulated gene expression is shown for reference (red, dashed, vertical line). P value shown for two-sided one sample t-test. **i)** Similar to (h), but selecting gene sets randomly from all cell-cycle genes annotated by KEGG^96^ or Gene Ontology^97^ (n = 1,801 cell cycle genes). **j)** Violin plots showing the distribution of DREAM activity values in individual samples (n = 1,101 individuals), Z-score normalized within each brain tissue, stratified by the duration each individual had AD before death. No pairwise comparisons between AD duration groups (violin plots) were significant (p < 0.05) based on two-sided Mann-Whitney U tests.

**Supplementary Figure 7:**
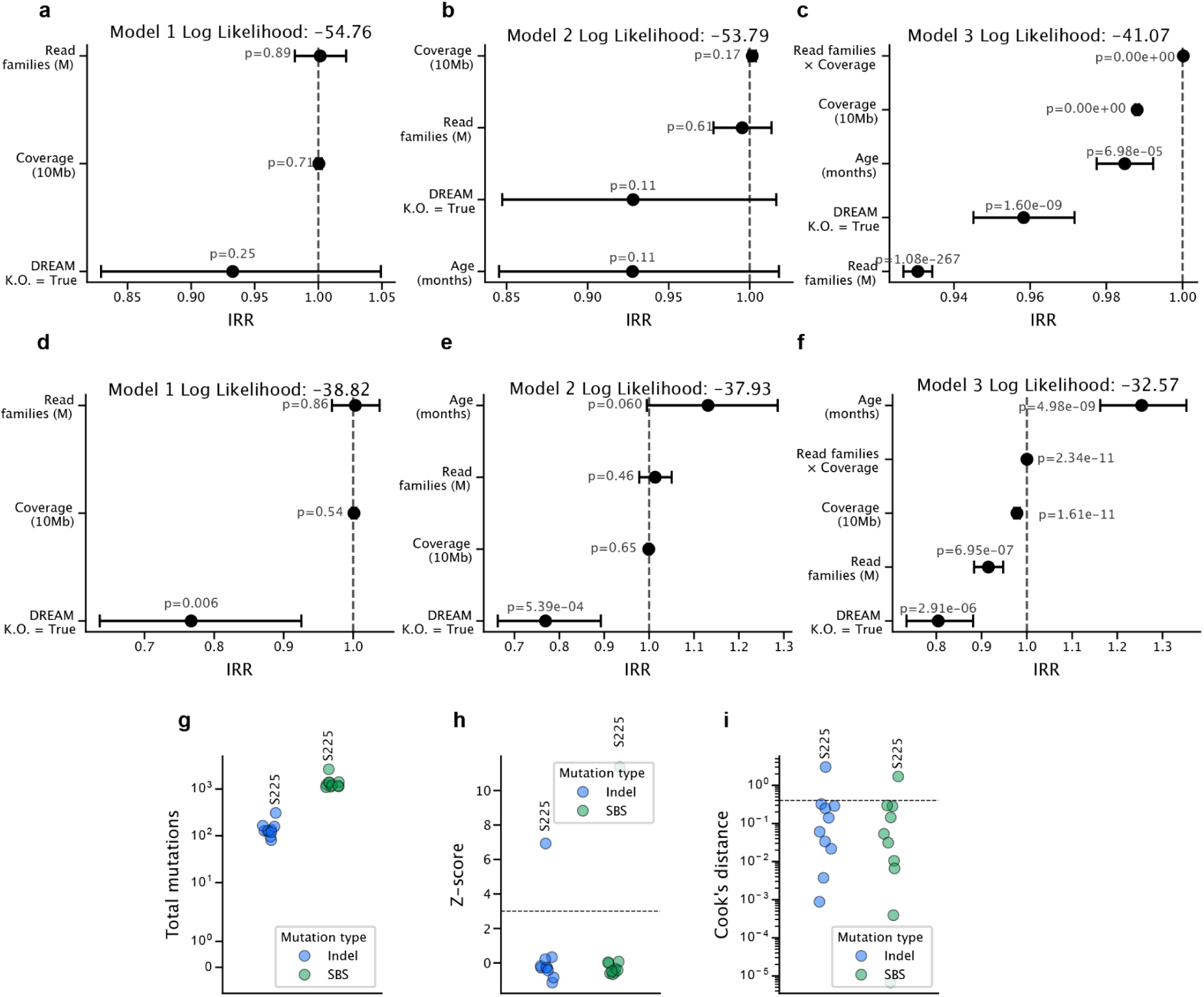
Duplex sequencing. **a-c)** Associations between covariates and single-base substitution (SBS) mutations per cell from negative binomial regression (Methods). Points and whiskers show incidence rate ratios (IRR) with 95% CIs. Model 1 (e) includes DREAM K.O. status, read families (per million) and coverage (per 10 Mb). Model 2 (f) adds age (months). Model 3 (g) further includes a read-families × coverage interaction. P values are from two-sided Wald tests. Less-negative log-likelihood indicates better fit. **d-f)** Similar to (a-c), but for insertion/deletion (ID) mutations. **g–i)** Per-mouse mutation burden and outlier diagnostics by mutation type. Each point is one mouse; colors denote mutation class, with outlier sample S225 labeled. (g) Total mutations per mouse (log_10_ scale). (h) Z-scores used to assess outlier status – the dashed line marks the Z-score threshold of 3. (i) Cook’s distance from the regression model (log_10_ scale) – the dashed line marks the Cook’s distance threshold of 1. Samples above the thresholds on all mutation classes are considered outliers.

