## Supplementary Figures and Tables for "The DREAM complex links somatic mutation, lifespan, and disease"

#### Supplementary Table 1

| | Description | Citation | Data type | Tissue | Pre-processing |
|---|---|---|---|---|---|
| 1 | Tabula Muris Senis | [33] | Smart-seq2 & somatic mutation identification | Pan-tissue | See “scRNA-seq processing” section |
| 2 | DREAM treated U2OS cells and *C. elegans* | [23] | RNA-seq | U2OS cells & *C. elegans* | See original publication |
| 3 | Cross-species transcriptomics | [35] | RNA-seq | Liver, kidney, and brain | See original publication |
| 4 | Cross-species somatic mutation rates | [4] | Somatic mutation rate quantification | Colon | See original publication |
| 5 | Cells treated with damaging agents | [2] | LD50 | Fibroblasts | See original publication |
| 6 | Quantification of single strand breaks & abasic sites in mouse tissues | [95] | SSiNGLe & SSiNGLe-AP assays | Multiple tissues | See original publication |
| 7 | Inbred mouse strains | [40] | Lifespan & RNA-seq | Liver | See original publication |
| 8 | Tabula Sapiens | [42] | 10X scRNA-seq | Pan-tissue | See “scRNA-seq processing” section |
| 9 | SEA-AD | [81] | 10X snRNA-seq & neuropathology | Brain | See “scRNA-seq processing” section |
| 10 | ROSMAP | [44] | RNA-seq & neuropathology | Brain | See original publication |

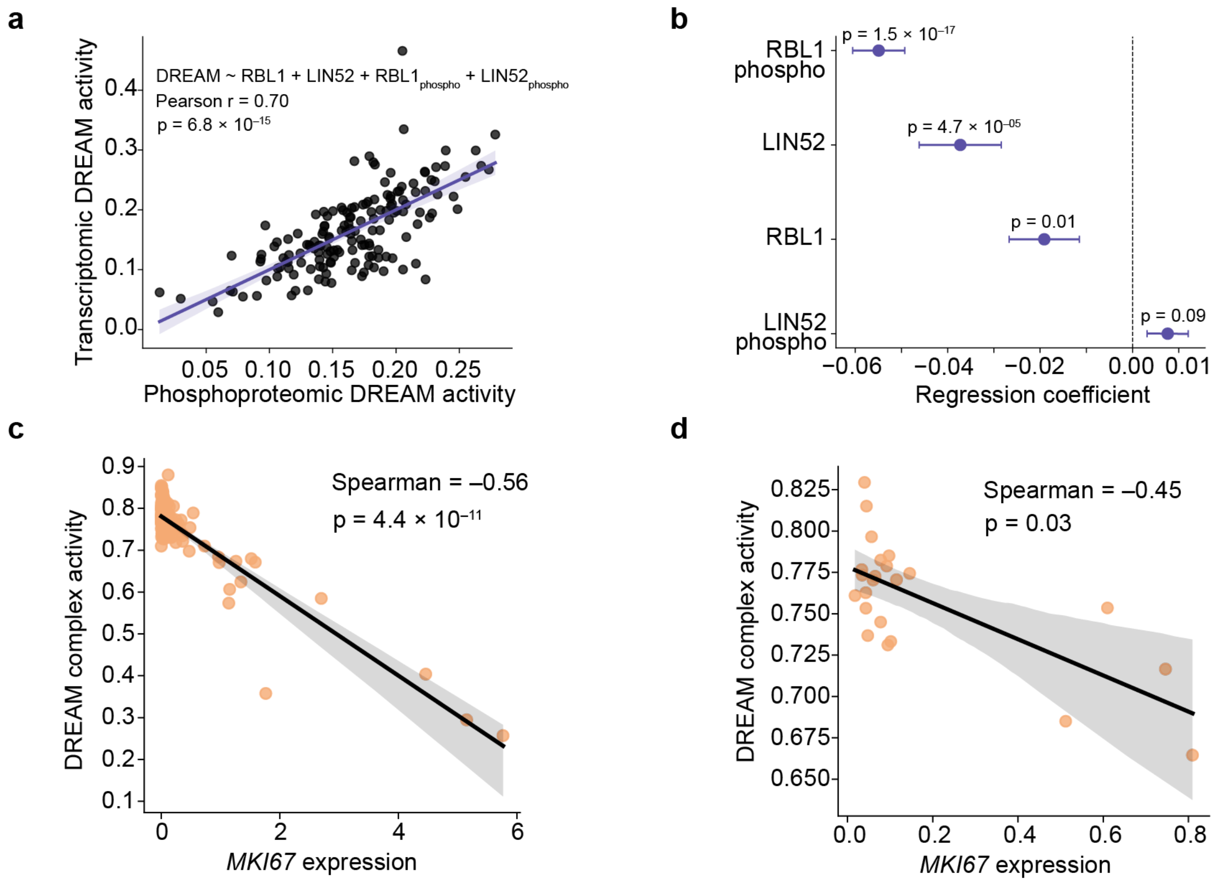


**Supplementary Figure 1: DREAM target expression mirrors other activity measures**

**a)** Scatterplot of DREAM activity in human tumor tissues inferred from either the protein abundance and phosphorylation status of DREAM complex components (x-axis) or the gene expression of DREAM-target genes ("transcriptomic DREAM activity", y-axis, n = 160 individuals). Two-sided p value calculated based on the exact distribution of Pearson's r modeled as a beta function.

**b)** Association of protein abundance and phosphorylation status of DREAM complex components (RBL1 and LIN52) with transcriptomic DREAM activity in human tissues (n = 160 individuals). Dots and whiskers indicate the coefficient and standard error from a linear mixed effects model regressing DREAM activity on the variables shown (**Methods**). Significance was calculated using two-sided t-tests.

**c)** Scatter plot of the mean expression of the proliferation marking gene MKI67 versus the mean DREAM complex activity, in each cell type (n = 120 cell types). Two-sided p value calculated modeling Spearman ρ's as a Student's t distribution.

**d)** Similar to (c) but in each tissue type (n = 21 tissues).

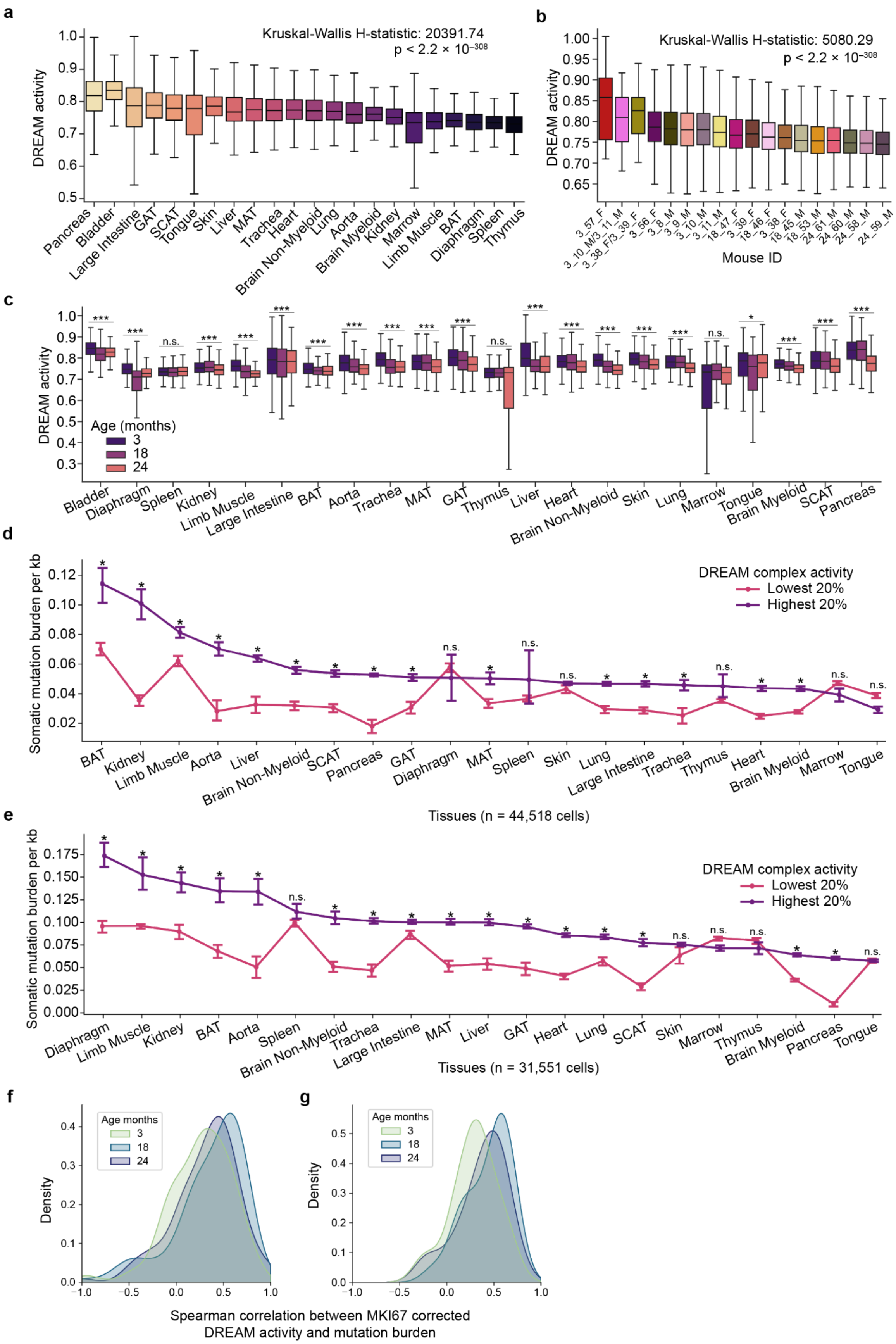
a
Kruskal-Wallis H-statistic: 20391.74
p < 2.2 × 10−308
DREAM activity
Pancreas
Bladder
Large Intestine
GAT
SCAT
Tongue
Skin
Liver
MAT
Trachea
Heart
Brain Non-Myeloid
Lung
Aorta
Brain Myeloid
Kidney
Marrow
Limb Muscle
BAT
Diaphragm
Spleen
Thymus
b
Kruskal-Wallis H-statistic: 5080.29
p < 2.2 × 10−308
DREAM activity
Mouse ID
c
DREAM activity
Age (months)
3
18
24
d
Somatic mutation burden per kb
DREAM complex activity
Lowest 20%
Highest 20%
Tissues (n = 44,518 cells)
e
Somatic mutation burden per kb
DREAM complex activity
Lowest 20%
Highest 20%
Tissues (n = 31,551 cells)
f
g
Age months
Density
Spearman correlation between MKI67 corrected
DREAM activity and mutation burden

**Supplementary Figure 2: DREAM activity in single cells**

**a)** Box plots of the distribution of DREAM complex activity by tissue across all cells from all mice (n = 110,824 cells, n = 18 mice). Two-sided p value from a Kruskal-Wallis test for a difference of distribution between tissues is shown. Boxes show inter-quartile range (IQR) with median line; whiskers extend to 1.5 IQR

**b)** Box plots of the distribution of DREAM complex activity by mouse across all cells from all mice (n = 110,824 cells, n = 18 mice). Two-sided p value from a Kruskal-Wallis test for a difference of distribution between tissues is shown.

**c)** Box plots of the distribution of DREAM activity scores in each tissue at each age (n = 110,824 cells, n = 18 mice). (***), (*), and (n.s.) indicate p values calculated by modeling Spearman ρ's as a Student's t distribution of $p < 1.0 \times 10^{-7}$, $p < 0.01$, and $p \geq 0.01$, respectively.

**d)** The somatic mutation burden per kilobase in cells (n = 44,518 cells, n = 10 mice) stratified by DREAM complex activity in 3 month old mice. Cells were partitioned into those with the highest and lowest 20% of DREAM complex activities within each tissue. Error bars denote 95% confidence intervals. (*) indicates $p < 0.0023$ (Bonferroni corrected p-value) and (n.s.) indicates $p \geq 0.0023$, based on a two-sided Mann-Whitney test.

**e)** Similar to (c), but for cells from 24 month old mice (n = 31,551 cells, n = 4 mice).

**f)** The distribution of Spearman correlation coefficients between MKI67-corrected DREAM complex activity and somatic mutation burden, within each cell type and age (n = 120 cell types, n = 110,824 cells, **Methods**).

**g)** Similar to (e) but within each tissue type (n = 21 tissues, n = 110,824 cells).

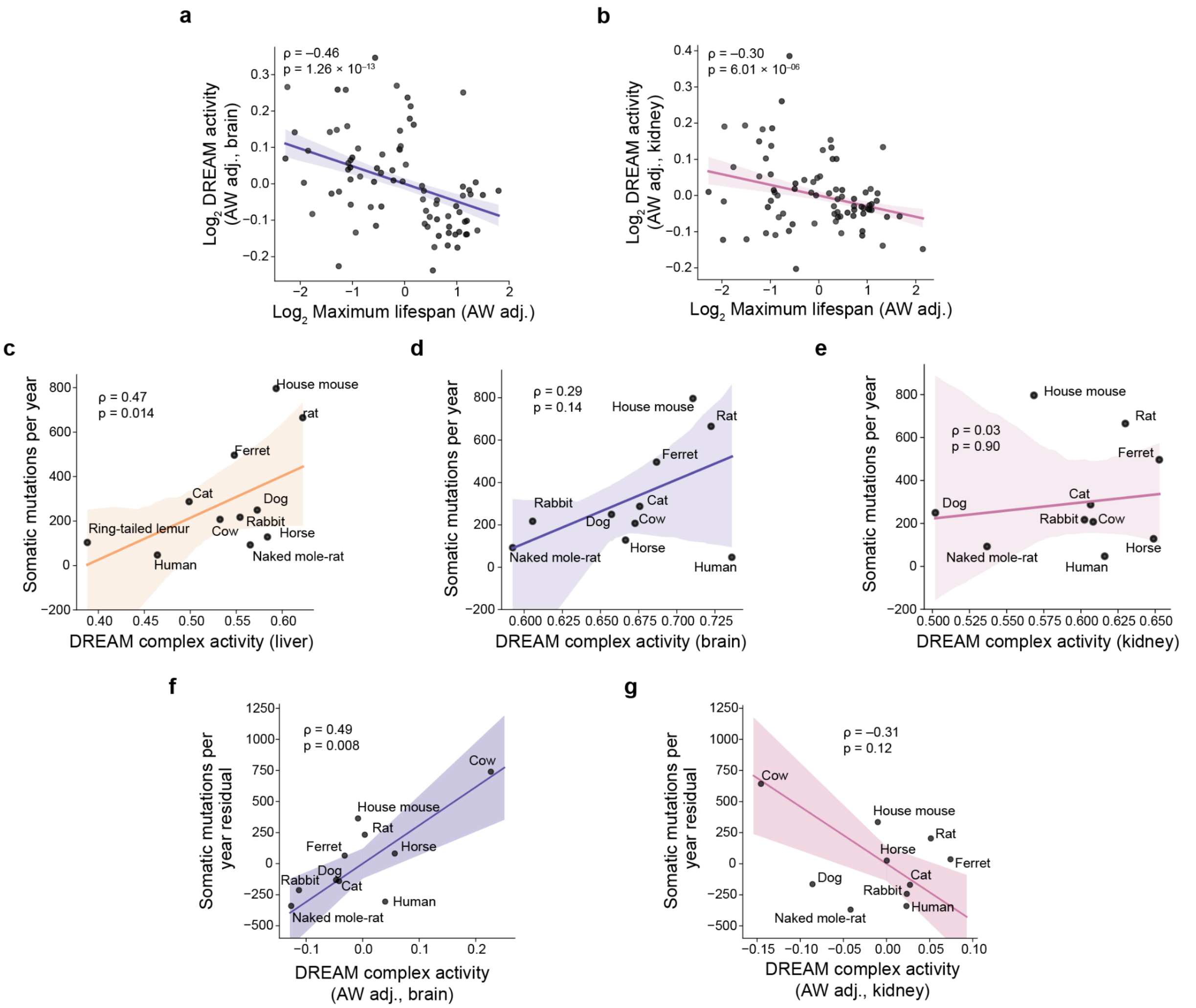


**Supplementary Figure 3: Mutation rate across species**

**a)** Scatterplot and reduced major axis regression depicting the average adult weight (AW) adjusted DREAM activity and lifespan across species (n = 92 species, **Methods**). Both axes are shown on a logarithmic scale.

**b)** Similar to (a), but DREAM activity is measured in the kidney.

**c)** Scatterplot and reduced major axis regression of the mean DREAM activity in the liver of a species and somatic mutation rate in the colon in that same species (n = 11 species, **Methods).**

**d)** Similar to (c), but DREAM activity is measured in the brain.

**e)** Similar to (c), but DREAM activity is measured in the kidney.

**f)** Scatterplot and reduced major axis regression depicting the average AW adjusted DREAM activity and somatic mutation rate (in the brain and colon, respectively) across species (n = 11 species, **Methods**).

**g)** Similar to (f), but DREAM activity measured in the kidney.

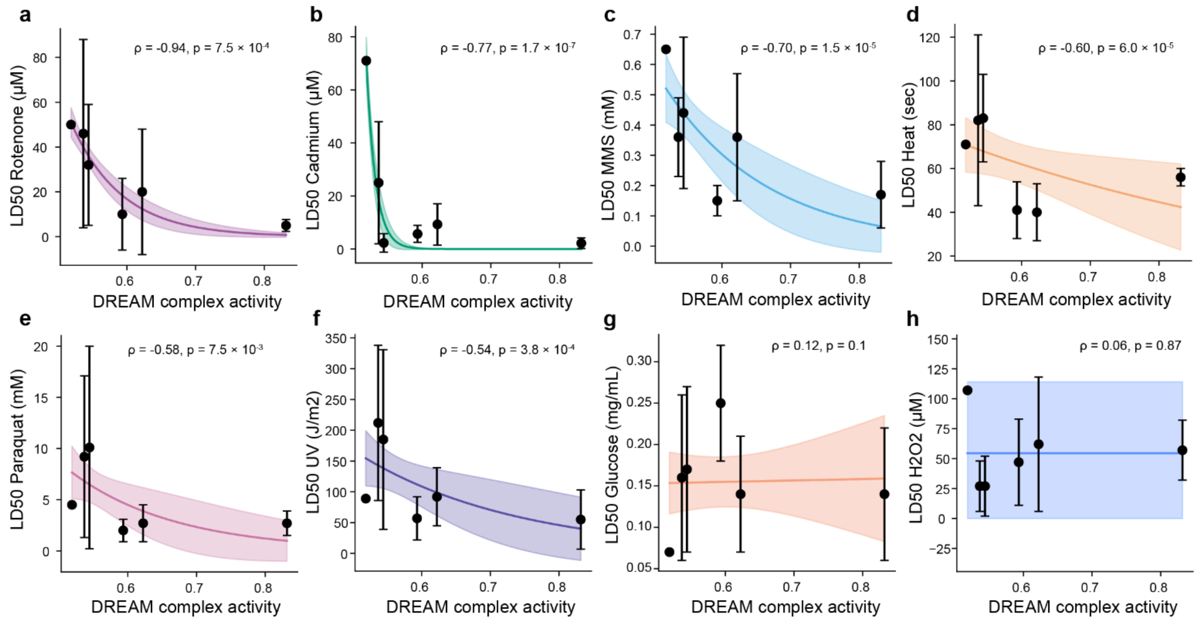


**Supplementary Figure 4: Connection of DREAM activity with DNA damage**

**a-h)** Scatter plots of species-specific liver DREAM complex activity vs. resistance (LD50) of fibroblasts from six species to damaging agents (from Harper et al.[2]). The points and error bars indicate the mean and 95% confidence interval of the LD50 of all cell lines of a species. The colored line and shaded area indicate the fit of an exponential decay model and 95% confidence interval, respectively. From left to right in each plot species include: the north american beaver (n = 1 cell line), red squirrel (n = 9 cell lines), white-footed mouse (n = 7 cell lines), house mouse (n = 9 cell lines), Norway rat (n = 6 cell lines), and deer mouse (n = 5 cell lines). Significance was calculated using a two-sided t-test between the cell lines of the 3 species with highest and lowest DREAM activity, respectively.

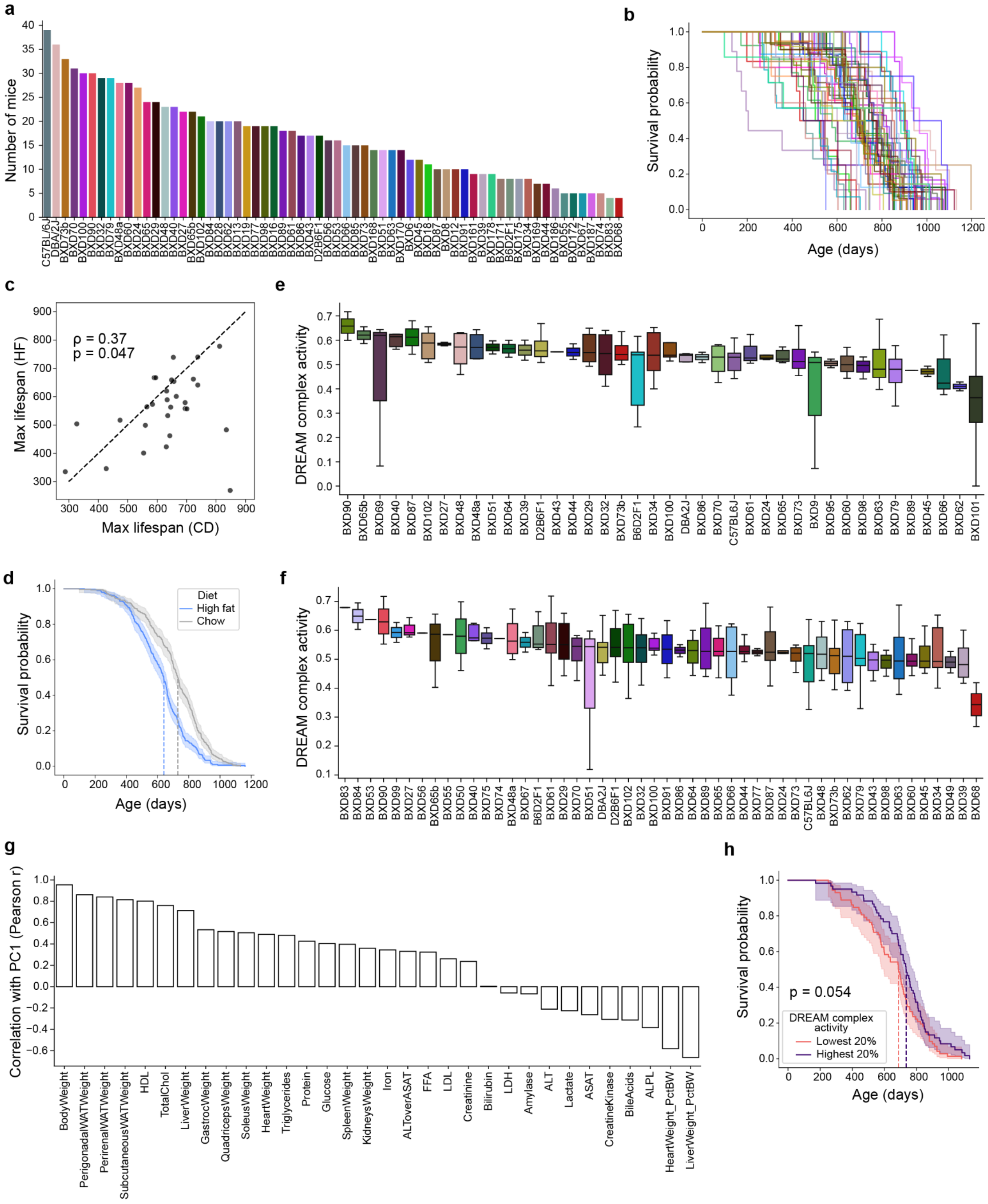

**Supplementary Figure 5: Survival and DREAM activity in mice**

**a)** Bar chart of the number of mice of each strain (n = 50 strains, n = 882 mice).

**b)** Kaplan-Meier survival curve for each strain in (a).

**c)** Scatterplot comparing the maximum lifespan of each strain on the chow diet (CD, x-axis) to the high-fat diet (HFD, y-axis, n = 29 strains profiled for lifespan and RNA-seq on both diets).

**d)** Kaplan-Meier survival curve for mice of all strains on the CD (n = 498 mice) compared to the HFD (n = 522 mice). The dark line and shaded area indicate the proportion of mice surviving to a particular age and the 95% confidence interval of this survival estimate, respectively. Dashed lines indicate the median survival of each group.

**e)** Box plots of the distribution of DREAM complex activity in mice of each strain fed the HFD (n = 120 mice profiled for liver gene expression).

**f)** Similar to (e) but for mice fed the CD (n = 150 mice profiled for liver gene expression).

**g)** Correlation of the standardized values of each covariate with principal component 1 (i.e., "PC1 loading", n = 270 mice, **Methods**).

**h)** Kaplan-Meier survival curve of CD fed mice belonging to strains with the highest vs lowest 20% of DREAM complex activities (purple vs. pink, n = 132 mice, n = 15 strains). The dark line and shaded area indicates the proportion of mice surviving to a particular age and the 95% confidence interval of this survival estimate, respectively. Dashed lines indicate the median survival of each group. P value calculated from a two-sided log-rank test.

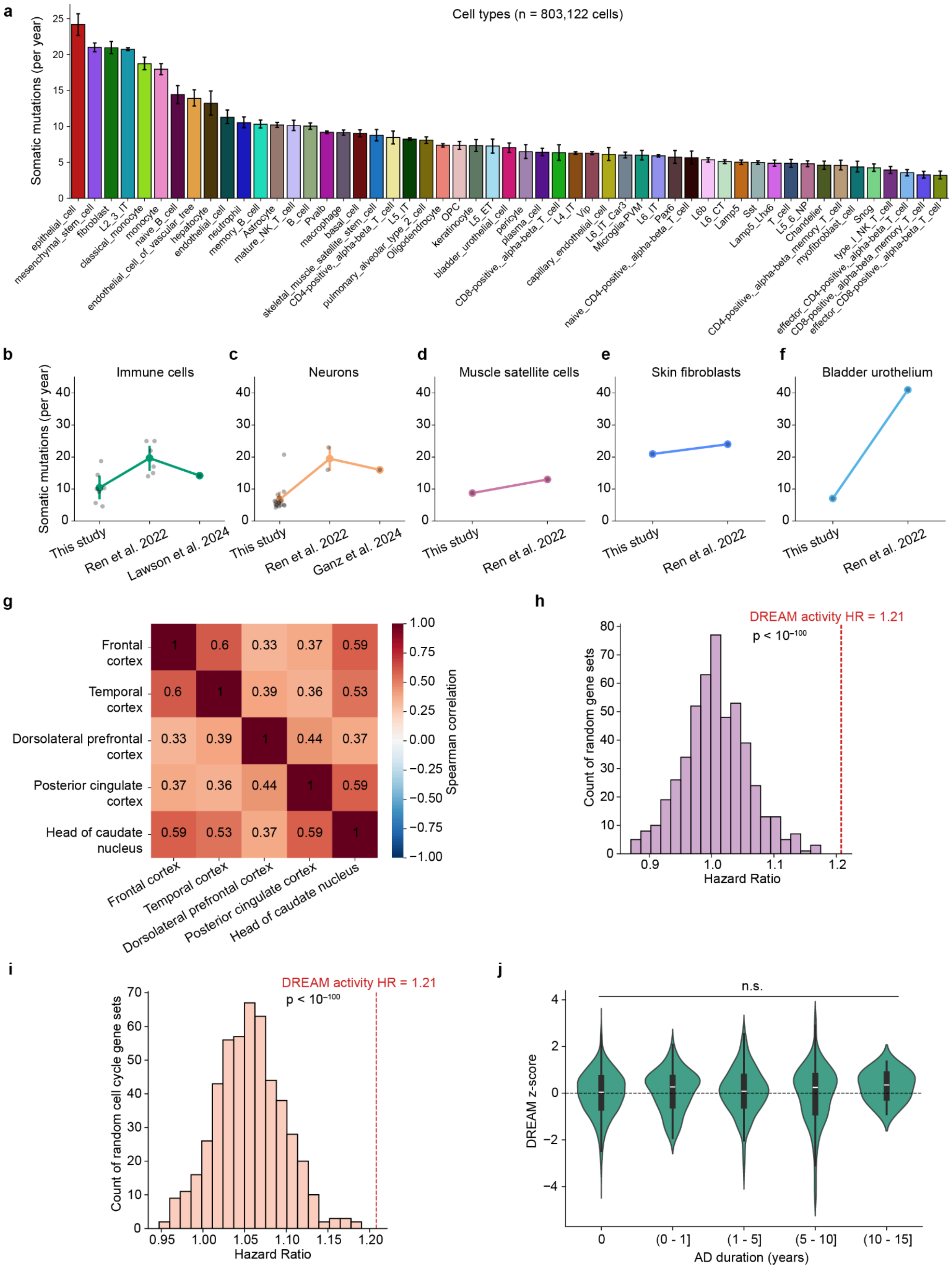
a
Cell types (n = 803,122 cells)
Somatic mutations (per year)
epithelial_cell
mesenchymal_stem_cell
fibroblast
L2_3_IT
classical_monocyte
monocyte
naive_B_cell
endothelial_cell_of_vascular_tree
hepatocyte
endothelial_cell
neutrophil
memory_B_cell
Astrocyte
mature_NK_T_cell
B_cell
Pvalb
macrophage
basal_cell
skeletal_muscle_satellite_stem_cell
CD4-positive,_alpha-beta_T_cell
L5_IT
pulmonary_alveolar_type_2_cell
Oligodendrocyte
OPC
keratinocyte
L5_ET
bladder_urothelial_cell
pericyte
plasma_cell
CD8-positive,_alpha-beta_T_cell
L4_IT
Vip
capillary_endothelial_cell
L6_IT_Car3
Microglia-PVM
L6_IT
Pax6
naive_CD4-positive,_alpha-beta_T_cell
L6b
L6_CT
Lamp5
Sst
Lamp5_Lhx6
T_cell
L5_6_NP
Chandelier
CD4-positive,_alpha-beta_memory_T_cell
myofibroblast_cell
Sncg
type_I_NK_T_cell
effector_CD4-positive,_alpha-beta_T_cell
CD8-positive,_alpha-beta_memory_T_cell
effector_CD8-positive,_alpha-beta_T_cell
b
Immune cells
Somatic mutations (per year)
This study
Ren et al. 2022
Lawson et al. 2024
c
Neurons
This study
Ren et al. 2022
Ganz et al. 2024
d
Muscle satellite cells
This study
Ren et al. 2022
e
Skin fibroblasts
This study
Ren et al. 2022
f
Bladder urothelium
This study
Ren et al. 2022
g
Frontal cortex
Temporal cortex
Dorsolateral prefrontal cortex
Posterior cingulate cortex
Head of caudate nucleus
Spearman correlation
h
DREAM activity HR = 1.21
p < 10^-100
Count of random gene sets
Hazard Ratio
i
DREAM activity HR = 1.21
p < 10^-100
Count of random cell cycle gene sets
Hazard Ratio
j
n.s.
DREAM z-score
AD duration (years)
0
(0 - 1]
(1 - 5]
(5 - 10]
(10 - 15]

**Supplementary Figure 6: Somatic mutations in human tissues**

**a)** Barplot indicating the mean and 95% confidence interval of the number of somatic mutations identified in each cell per year (normalized to haploid genome size, **Methods**) grouped by cell type (n = 803,122 cells, n = 90 individuals).

**b-f)** Line plots comparing the number of somatic mutations per year identified by "this study" compared with previous estimates using DNA sequencing. Dark points indicate the mean mutation rate for a single cell type in a particular study, while the colored points indicate the mean and 95% confidence interval of these mutation rates. The x-axis lists the study from which the respective data came.

**g)** Heatmap depicting the pairwise Spearman correlations of DREAM complex activity in different brain regions of the same individual (n = 1,101 individuals, 2,776 samples, mean 2.4 ± std 1.0 brain regions per individual). All pairwise correlations are significant ($p < 0.0004$).

**h)** Histogram showing the distribution of hazard ratios from Cox proportional hazards models predicting age at AD diagnosis (using the same covariates as **Fig. 6g**, n = 1,101 individuals, **Methods**) using the ssGSEA enrichment of 500 randomly chosen sets of genes. HR of DREAM-regulated gene expression is shown for reference (red, dashed, vertical line). P value shown for two-sided one sample t-test.

**i)** Similar to (h), but selecting gene sets randomly from all cell-cycle genes annotated by KEGG[96] or Gene Ontology[97] (n = 1,801 cell cycle genes).

**j)** Violin plots showing the distribution of DREAM activity values in individual samples (n = 1,101 individuals), Z-score normalized within each brain tissue, stratified by the duration each individual had AD before death. No pairwise comparisons between AD duration groups (violin plots) were significant ($p < 0.05$) based on two-sided Mann-Whitney U tests.

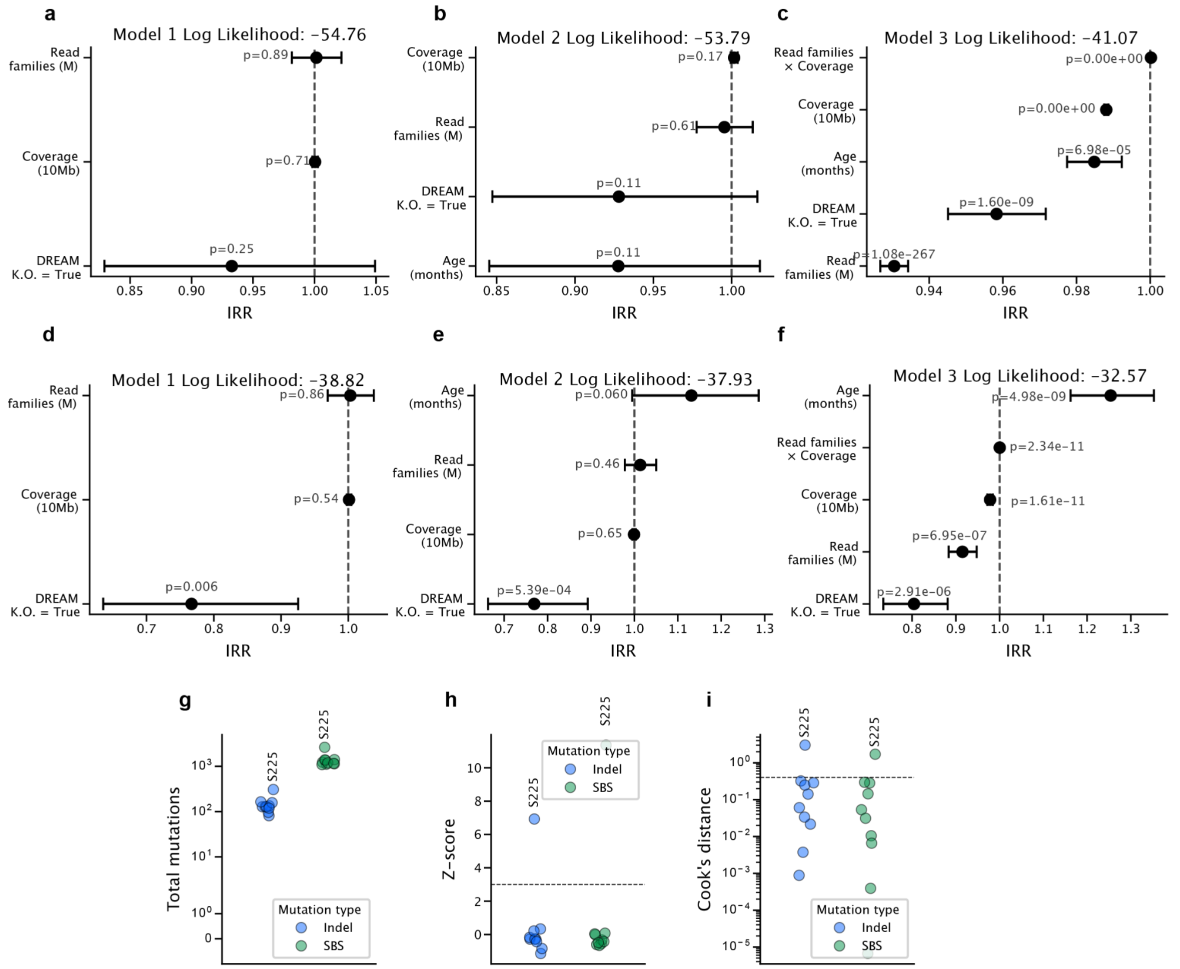


### Supplementary Figure 7: Duplex sequencing

**a-c)** Associations between covariates and single-base substitution (SBS) mutations per cell from negative binomial regression (Methods). Points and whiskers show incidence rate ratios (IRR) with 95% CIs. Model 1 (e) includes DREAM K.O. status, read families (per million) and coverage (per 10 Mb). Model 2 (f) adds age (months). Model 3 (g) further includes a read-families × coverage interaction. P values are from two-sided Wald tests. Less-negative log-likelihood indicates better fit.

**d-f)** Similar to (a-c), but for insertion/deletion (ID) mutations.

**g–i)** Per-mouse mutation burden and outlier diagnostics by mutation type. Each point is one mouse; colors denote mutation class, with outlier sample S225 labeled. (g) Total mutations per mouse ($log_{10}$ scale). (h) Z-scores used to assess outlier status – the dashed line marks the Z-score threshold of 3. (i) Cook's distance from the regression model ($log_{10}$ scale) – the dashed line marks the Cook's distance threshold of 1. Samples above the thresholds on all mutation classes are considered outliers.
